# Leukemia stem cell expansion cultures reveal clonal drivers of leukemogenesis and therapy response

**DOI:** 10.64898/2026.02.24.707683

**Authors:** Indranil Singh, Andrea Polazzi, Alex Maya Pombo, Marta López-Osias, Celine Bauer, Maria Guarini, Pedro Sánchez Sánchez, Luna Goulet, Carmen Gallardo, Daniel Fernández-Perez, Robert L. Bowman, Alejo E. Rodriguez-Fraticelli

## Abstract

Leukemia stem cells (LSCs) contain the highest capacity for leukemia-reinitiation and therapy-resistance across all leukemic cells, but our understanding of their molecular and cellular properties remains limited due to their relative rarity and ineffective cell culture systems to maintain their purity at scale. Here, we develop Polymer-based Leukemic STem-cell Cultures (PLSTCs) and demonstrate their capacity to derive and propagate large numbers of Npm1^cA^/Flt3^ITD^ acute myeloid leukemia (AML) stem cells at high purities. Compared to traditional cultures, PLSTCs show more than 1000-fold enrichment in functional LSCs based on single-cell gene expression signatures and leukemia-initiating assays. Tracing LSC clones with genomic LARRY barcodes during *ex vivo* expansion, we reveal that PLSTCs can sustain a diversity of self-renewing LSC states with stable, heritable transcriptional programs. Using dynamic state-fate analysis, we characterize clonal programs that are linked with enhanced *ex vivo* self-renewal, *in vivo* leukemia initiation, and therapeutic response to induction chemotherapy. LSC clones primed to resist treatment were enriched for a rare cell state that underwent a fate-switch and produced megakaryocytic-erythroid-like leukemic cells that expanded in the spleen. Targeting LSC programs through pooled CRISPR and single-cell sequencing (CROPseq) in PLSTCs, we reveal that chondroitin-sulfate synthesis is required to maintain a primitive LSC state and leukemic recovery from chemotherapy. In sum, our studies showcase the powerful application of scalable leukemic stem-cell expansion cultures and dynamic state-fate analysis of AML LSCs. We anticipate these systems will accelerate our understanding and interception of stem cell plasticity in cancer.

## Introduction

Cellular plasticity drives cancer evolution by enabling transitions between heterogeneous phenotypic states^1–4^. In acute myeloid leukemia (AML), the key drivers of plasticity are thought to act within leukemic stem cells (LSCs), the rare cells that are capable of sustaining disease, fueling relapse, and resisting therapy^5–9^. However, studying the clonal trajectories and regulatory programs in LSCs remains a challenge due to the unreliability of markers and their rarity both *in vitro* and *in vivo*.

Phenotypic hierarchies in AML are also strikingly heterogeneous patient to patient, and within patients, particularly within the most frequent *de novo* AML, which carry the NPM1 mutation^10–14^. However, it is unclear how much of this variation reflects intrinsic, stable, and heritable differences versus extrinsic influences and short-term fluctuations. Disentangling these sources of heterogeneity requires studying LSCs in a controlled, uniform environment and tracking how individual cells maintain or alter their transcriptional states^15–17^.

Current *ex vivo* culture systems of primary AML LSCs can maintain them short-term, and at extremely low frequencies, and this is evidenced by the large numbers of cells that are necessary to transplant to reinitiate leukemias^18–20^. Expansion of LSCs is particularly difficult for NPM1^cA^/FLT3^ITD^-mutant AMLs, the most frequent aggressive adult subtype. To define the intrinsic logic of leukemic stemness and plasticity, a system is needed that supports sustained, efficient LSC propagation to track their mechanisms of heterogeneity and plasticity.

Polymer-based culture systems have revolutionized our capacity to expand and study self-renewing, transplantable mouse and human HSCs^21–26^. Building on these innovations, we develop PLSTCs, Polymer-based Leukemic STem-cell Cultures. We show that PLSTC can propagate mouse Npm1^cA^/Flt3^ITD^ AML LSCs at high purity (at least 100-fold enrichment over conventional Stemspan cultures). Tracing LSC clonal heterogeneity *ex vivo*, we reveal heritable transcriptional programs coupled to differences in *ex vivo* self-renewal, *in vivo* leukemia initiation, and chemotherapy resistance. Finally, we use PLSTCs to carry out a targeted perturbation screen using single-cell read-out (CROPseq), identifying *Csgalnact1* as a regulator of the resistant-primed, self-renewing LSC program. Together, PLSTCs provide a scalable experimental foundation for mechanistic and translational interrogation of primary AML LSCs.

### PLSTCs establish an LSC-enriched, scalable platform that preserves stemness and reveals functional differences to conventional culture

To test whether defined, bovine serum albumin (BSA)-free conditions preserve stemness and functional properties of LSCs ex vivo, we isolated and cultured mouse bone marrow (BM) AML cells derived from a recently developed Flp-inducible Npm1^cA^/Flt3^ITD^ AML model (see Methods)^27^. AML cells were cultured for 30 days in Stemspan or PLSTC medium, supplemented with 10 ng/ml IL3, 10 ng/ml IL6, 50 ng/ml SCF, and 100 ng/ml TPO (**Figure 1a**; see Methods for culture system details)^27^. While both expansion cultures supported proliferation and long-term expansion of leukemic cells, compared to Stemspan cultures, PLSTCs had a 40% reduced doubling rate (0.56 doublings per day, **Figure 1b**). Phenotypically, PLSTC cultures yielded smaller-sized cells, with a pronounced shift towards a CD34^+^ CD16/32^−^ (FcgammaR^−^) immature-like compartment (ranging from 51% to 78% of all cells) while Stemspan cultures maintained a lower fraction of this putative LSC-like compartment^9,28^ (**Figure 1c**). These results suggest an enrichment of primitive LSC-like cells in the PLSTC conditions.

**Figure 1.**
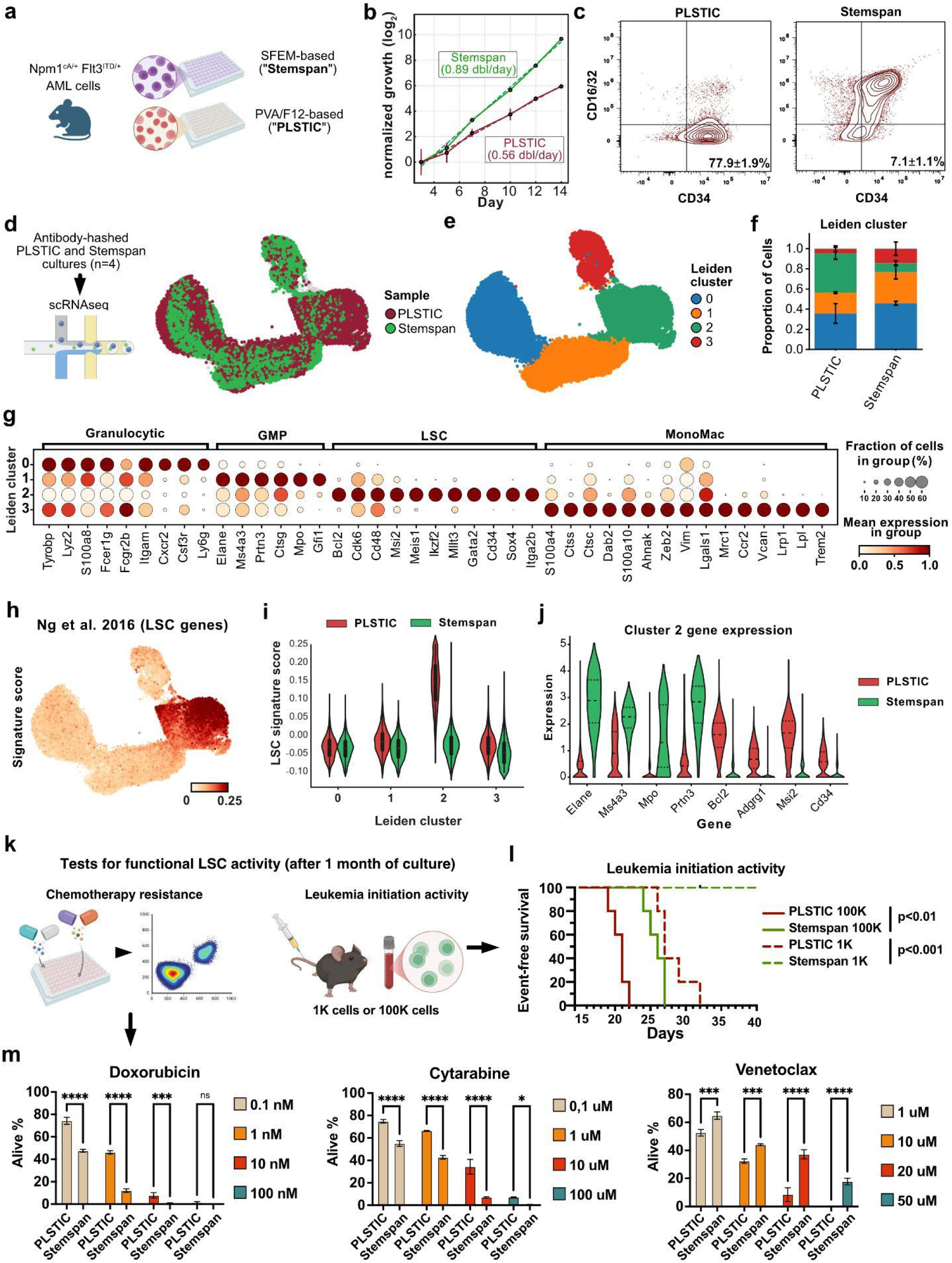
PLSTCs establish an LSC-enriched, scalable platform that preserves stemness and reveals functional differences to conventional culture. **a**) Npm1^cA^ Flt3^ITD^ AMLs were established by transplanting Npm1^frt-stop-cA^ Flt3^frt-stop-ITD^ LSKs transduced with a Flpo-EGFP lentivirus into lethally irradiated mice. Then, 10^5^ EGFP+ cells were cultured in Stemspan or PLSTC conditions for 30 days, and then assayed in various experiments. **b)** Normalized growth curves of Stemspan and PLSTC cultures using a log_2_ scale, showing the difference in doubling rates (n=2 independent experiments). **c)** Flow cytometry profiles of Stemspan and PLSTC cultures using the GMP/Myeloid differentiation marker CD16/32 (Fc-gamma receptors) and the LSC marker CD34. **d)** Single-cell profiling and UMAP plots comparing Stemspan and PLSTC cultures (n=4 cultures). **e)** Leiden clustering of Stemspan and PLSTC cultures reveals 4 major clusters. **f)** Differential proportion analysis showing PLSTC cultures enriched in cluster 2. **g)** Dotplot showing selected top markers of various clusters, highlighting the specificity of LSC markers in cluster 2. **h)** Human CD34+/CD38- LSC signature score (Ng et al. 2017) in the *ex vivo* LSC culture UMAP. **i)** Violin plot showing the specific expression of the LSC signature score in PLSTC LSCs within cluster 2. **j)** Violin plot showing different LSC and Maturation markers for each culture condition. **k)** Functional tests of LSC phenotypic activity. **l)** Survival curves at different transplantation cell doses comparing PLSTC and Stemspan cultures (n=5 transplanted mice). **m)** Dose-response efficacy of various anti-leukemic therapeutic compounds targeting cell-cycle (cytarabine, araC, and doxorubicin, Doxo) or metabolic targeting through Bcl2 (venetoclax) (n=4 independent cultures). Two-sided t-tests *p<0.05, **p<0.01, ***p<0.005, ****p<0.001

Next, to resolve cell states at single-cell resolution, we pooled TotalSeq-hashed cell culture expansion experiments (n = 4) and performed scRNA-seq (**Figure 1d**). Unsupervised analysis identified four major Leiden clusters (**Figure 1e**), with PLSTC cells being significantly enriched in cluster 2 (**Figure 1f**), which expressed the highest levels of leukemia stem cell markers, including *Cd34* (CD34), with a 5.1 fold-change (adj. P = 10^−300^), *Adgrg1* (GPR56, 5.3 fold-change, adj. P = 10^−300^), and *Msi2* (Musashi-2, 4.5 fold-change, adj. P < 10^−300^)^29,30,11^ (**Figure 1g and Supplementary Table 1**). In contrast, Stemspan cultures were enriched in cell-states that annotated to a granulocytic/neutrophil-like program (marked by expression of *Ly6g*, *Cxcr2, Fcer1g*)^31^, a GMP-like compartment (*Mpo, Prtn3*, *Elane*), or a monocytic/macrophage-like program (*Ccr2*, *Trem2*, *Ctss*)^32^, confirming the inefficacy of conventional culture systems to maintain undifferentiated cells at high frequency.

We next calculated an LSC score using published human datasets of CD34+/CD38- LSCs (**Figure 1h and Extended Data Figure 1d-e**)^33^. The LSC gene signature revealed a specific enrichment in cluster 2, but only in PLSTC cells, which also showed higher expression levels of various LSC marker genes compared to Stemspan cells in that same cluster (**Figure 1i-j, Extended Data Figure 1a, and Supplementary Table 2**). This indicated that, while cluster 2 marks the most primitive cells in both culture conditions, only PLSTC cultures can expand and support primary LSC transcriptional programs that resemble human LSC gene expression.

We next asked whether this transcriptional enrichment translated into functional LSC activity. We established two orthogonal assays (**Figure 1k**). First, we performed tail-vein AML cell transplantation into sub-lethally irradiated (900 cGy in split dosing) recipients at limiting cell doses (100, 1000, 100,000 whole culture cells) to read out tumour-initiating activity *in vivo* (**see Methods**)^27,34^. PLSTCs showed accelerated leukemia onset relative to Stemspan when transplanting a conventional dose of 10^5^ cells, and they were also capable of rapidly initiating leukemias even at the low dose of 10^3^ cells (log-rank p < 0.001 for each comparison; **Figure 1l**). In contrast, Stemspan cultures could not generate leukemias at the lower doses (even after 6 months post-transplantation), demonstrating a significantly expanded (at least 100-fold) LSC frequency and fitness in PLSTC-expanded LSCs.

Then, we performed drug-based cell-killing assays for different therapeutic compounds (**Figure 1m**). PLSTC cultures were up to 4.5-fold more resistant to standard chemotherapeutic agents, in line with their reduced cycling rate, and reduced expression of cycling markers (*Mki67*) compared to StemSpan cultures (**Figure 1m, left and central panels, and Extended Data Figure 1f**)^10^. Next, we hypothesized that PLSTC cultures would be more sensitive to anti-BCL2 therapeutics (Venetoclax), which have been demonstrated to effectively kill primitive-like AML LSCs on the basis of their cell-type enriched BCL2 expression and specific amino-acid metabolism dependencies^35^. This was indeed the case, with Stemspan cultures being up to 4-fold more resistant, in agreement with the differences in Bcl2 gene expression levels across both culture conditions (2.05 fold-change)(**Figure 1m, right panel, and Extended Data Figure 1f**).

Together, these results demonstrate that PLSTCs create a scalable culture environment that stabilizes LSC programs and functional potential, at very high frequency, while maintaining long-term proliferative capacity. Furthermore, the rapid adaptation to culture conditions provides strong evidence that strikingly different states and fates can emerge mostly through non-genetic effects.

### Single-cell lineage tracing in LSCs uncovers a diversity of stable, heritable genetic programs

Hematopoietic stem cells show long-lasting, heritable programs that regulate their functional biases, which can be uncovered through state-fate clonal analysis and single-cell lineage tracing^17,36–41^. To uncover similar mechanisms underlying differentiation and clonal output, we used lentiviral barcoding to trace *Npm1^cA^ Flt3*^ITD^ LSCs from their origins for up to 20 days in PLSTC culture. First, we isolated non-leukemic HSCs from *Rosa26-Flpo^ERT2^*, *Npm1^frt-cA^ Flt3*^frt−ITD^ mice, transduced them with differently indexed EGFP LARRY libraries, and cultured them on PLSTC media containing tamoxifen (4OHT), to induce the expression of the AML founder mutations^27,36^. Then, at day 20 post-culture, we profiled half of the cultures by single-cell RNA sequencing, amplifying the LARRY barcode to group cells according to their clone of origin^36^. The remaining cells were split between *in vivo* AML reinitiation experiments, or long-term culture until day 40 in multiple separate wells, whereafter these were also analyzed by FACS-sorting, single-cell RNA sequencing, and barcode quantification (**Figure 2a**). With this design we were able to compare sister-cells among LSC clones and longitudinal tracking of clonal fate *in vivo* and *ex vivo* while controlling for confounders from extrinsic cues.

**Figure 2.**
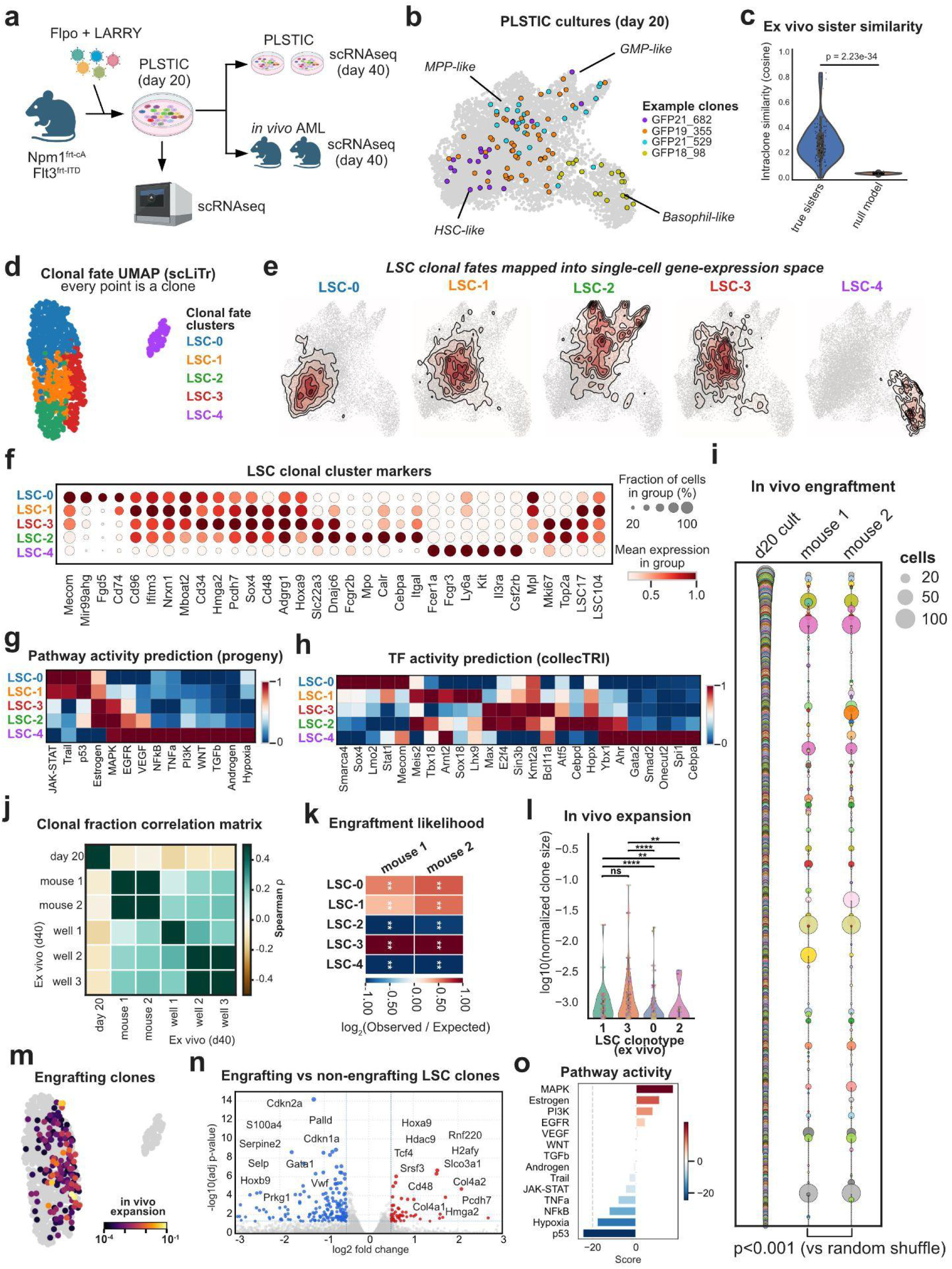
Clonal barcoding in PLSTCs uncovers discrete, heritable LSC programs. **a**) Npm1^frt-stop-cA^ Flt3^frt-stop-ITD^ HSCs were transduced with a barcoding LARRY library and Flpo-EGFP lentivirus and cultured for 20 days in PLSTC medium. Then, half of the EGFP+ cells were were profiled by scRNAseq (day 20), and the remaining cells were split into two assays: a long-term culture assay in PLSTC media (until day 40), and an *in vivo* AML initiation assay by transplantation (together with normal support BM) into lethally-irradiated mice (for 20 days), both of which sampled by scRNAseq at the end point. For all samples, single-cell cDNA libraries were used to prepare LARRY barcode-specific libraries by PCR and sequenced jointly to analyze clonal lineages. **b)** UMAP of PLSTC culture states at day 20, highlighting 4 example clones that occupy distinct territories of the UMAP, enriched in gene expression markers that correspond to the labeled categories. **c)** Violin plot showing the transcriptional memory within the clones, measured as sister cell gene expression similarity using the cosine distance in principal-component space, compared to a null model generated by randomly shuffling identities. **d)** Clonal fate UMAP computed using scLiTr. Every point represents a clone, and clones are grouped into fate clusters based on the distribution of their cells in the gene-expression UMAP space (fate-behavior similarity). **e)** Representation of fate clusters using density plots in the UMAP single-cell gene expression space. A different density plot is shown for each fate cluster (LSC-0 to LSC-4). **f)** Dotplot with transcriptional markers of distinct LSC fate clusters. Cell cycle markers *Mki67* and *Top2a*, and LSC gene scores are also shown in the right columns. **g)** Heatmap showing the predicted Progeny pathway activity score (based on univariate linear models) across different LSC fate clusters. **h)** Heatmap showing the predicted CollecTRI transcription factor activity score (based on univariate linear models) across different LSC fate clusters. **i)** Bubbleplot showing the size of clones in cell culture and *in vivo* initiated AMLs in separate recipient mice. Spearman correlation rho = 0.7, p<0.001. **j)** Correlation matrix of LARRY clone sizes comparing the initial cell culture (d20), the transplanted *in vivo* AMLs, and the long-term PLSTC cultures (d40). **k)** Heatmap showing the *in vivo* engraftment likelihood (observed vs expected over-representation analysis) comparing the different LSC fate clusters (adjusted for cell numbers). **l)** Violin plot showing the difference in *in vivo* clonal growth (fitness) comparing the LSCs based on their different initial fate cluster. **m)** Clonal fate UMAP colored by *in vivo* expansion LARRY clone sizes. Gray clones did not engraft. **n)** Volcano plot showing LSC differential gene expression results comparing *in vivo*-detected (engrafting) versus non-detected (non-engrafting) clones. **o)** Barplot showing the predicted Progeny pathway activity score in engrafting versus non-engrafting LSCs. Wilcoxon tests (corrected for multiple-tests) *p<0.05, **p<0.01, ***p<0.005, ****p<0.001

We detected 1375 clones (with 3 or more cells), with a median clone size of 6 cells (maximum size of 137 cells). Surprisingly, after 20 days of cell culture, involving 2 passages, and even splitting into separate wells, clonal cells were highly homogeneous, occupying a distinct, restricted subset of the entire transcriptional continuum (**Figure 2b, Extended Data Figure 2a**). Consequently, we verified that true sister cells were significantly more similar compared with random cell pairs (cosine similarity median of true sisters = 0.24, random sisters = 0.01, Wilcoxon test)(**Figure 2c**). This clonal homogeneity suggests that the phenotypic space of every clone is constrained, and thus, the observed transcriptional continuum is a composite of hundreds of restricted clonal manifolds.

To better classify these different self-renewing LSC states, we used scLiTr^42^, an analytic pipeline that groups clones based on their similarities of fate (i.e. its distributions across the state manifold), generating a clonal fate graph (**Figure 2d**). To use scLiTr, we first grouped single cell transcriptomes into Leiden clusters based on the gene expression manifold (**Extended Data Figure 2b-c**). Then, for each barcode-defined clone, we then computed the fraction of its cells in each Leiden cluster. Using this vector of fate fractions, and imputing putative dropouts, scLiTr assembled a clone-by-clone k-nearest neighbor graph. Applying clustering to the clonal graph itself yielded clonal fate clusters, which we term “LSC clonotypes”. This procedure groups clones by behavioral similarity rather than by any single marker and serves as our operational definition of LSC clonal programs.

Four major LSC clonotypes emerged *ex vivo* (underscored “*ev”*): LSC-0_ev_, LSC-1_ev_, LSC-2_ev_, and LSC-3_ev_. When projected back onto the cellular UMAP, these clonotypes localize to recognizable compartments defined by markers commonly observed in hematopoietic stem cells (HSC) and multipotent progenitor cells (MPPs) (**Figure 2e-f - Supplementary Table 3 and 4**)^32,43,44^. LSC-0_ev_ mapped to an HSC-like region with high expression of markers of normal long-term HSCs (*Fgd5*, *Mecom*)^45–47^. LSC-1_ev_ and LSC-3_ev_ were both enriched in markers of normal multipotent progenitors (*Cd34*, *Cd48, Sox4*)^48,49^, but LSC-3_ev_ showed higher expression of previously-reported LSC genes (*Adgrg1*/GPR56, *Hmga2*, *Hoxa9*)^50–52^. LSC-2_ev_ was enriched in GMP markers (*Mpo*, *Fcgr2b*), suggesting a more differentiated GMP-like LSC state^53^. An additional clonotype (fate cluster LSC-4_ev_) aligned with a granulocytic, mast-cell-like program (*Fcer1a*, *Csf2rb, Kit*). Thus, the different LSC clonotypes capture a range of self-renewing LSC states along a myeloid differentiation continuum, perhaps resembling the LSC heterogeneity observed in human AML samples.

To disentangle potential regulatory factors underpinning the stable differences in LSC programs, we used Decoupler^54^ to calculate transcription factor and pathway activity scores using a univariate linear model (ULM)-based on publicly-available network databases (CollecTRI, Progeny)^55,56^. These results revealed a similarity between LSC-0_ev_ and LSC-1_ev_ based on pathway activities (upregulated JAK-STAT, Trail, and p53 pathways), and a similar pathway convergence for LSC-2_ev_ and LSC-3_ev_ (enriched Estrogen, MAPK, and EGFR pathway scores) (**Figure 2g**). In contrast, transcription factor activity predictions were more specific at each end of the spectrum, with LSC-0_ev_ showing the highest scores for transcription factors associated with self-renewal, lymphopoiesis, and HSC emergence (*Lmo2*, *Sox4*)^57–59^, while LSC-2_ev_ and LSC-4_ev_ showed the highest scores for myeloid maturation regulators (*Cebpd, Ybx1, Gata2, Spi1, Cebpa*)^60^ (**Figure 2h**). The intermediate MPP-like clonotypes, LSC-1_ev_ and LSC-3_ev_, were principally enriched in pro-tumorigenic transcription factors (*Myc, Max, E2f4, Kmt2a*)^61^, suggesting a higher potential for continued leukemic fitness.

We next asked whether these *ex vivo* clonotypes were associated with different functional behaviors *in vivo*. Recipient animals transplanted with barcoded day-20 PLSTC LSCs generated *in vivo* leukemias with the expected latency (for 10,000 cells per animal), a median of 19 days post-transplantation (**Extended Data Figure 2d-f**). As expected, at the end point, the mice had enlarged spleens, leukemic bone marrow, and histopathological confirmation of blasts infiltrating lungs, spleens and liver. Complete blood counts of peripheral blood showed a staggering myelomonocytosis (Neutrophil and Monocyte counts > 50×10^3^/ul), lymphopenia, anemia, and thrombocytopenia (Lymphocyte counts < 2×10^3^/ul, Red blood cell counts < 5×10^6^/ul, Platelet counts < 500×10^3^/ul). Next, we sorted bone marrow LARRY-barcode+ (EGFP+) cells, and we sequenced 10,272 high-quality cells and their corresponding LARRY barcodes (see Methods). We detected 160 clones (with at least 3 cells per clone) across both recipients. To compare *ex vivo* and *in vivo* clone sizes for the same set of clones, we visualized the clone size distribution using a ranked bubble plot (**Figure 2i**). Clone sizes were highly correlating across each transplanted recipient mouse, confirming the influence of clone intrinsic factors in leukemic fitness^17^. We next compared *ex vivo* and *in vivo* outcomes by calculating a Spearman correlation matrix with all the LARRY clone size data across samples and replicates (**Figure 2j**). Within-assay correlations were always high (average Spearman Rho_mice_ = 0.56, average Rho_wells_ = 0.37). Correlations across assays showed a more modest, but still positive association (average Rho_mice-wells_ = 0.24), while comparisons to the initial clone size distribution revealed no association (average Rho_d20-d40_ = 0.06). These results suggest that *ex vivo* self-renewal and *in vivo* fitness properties are only partly linked, yet both are driven by a rare, deterministic subset of all the initial clones.

To define the molecular basis of these diverse functional properties, we first calculated which *ex vivo* clonotypes were more likely to engraft and reinitiate leukemias *in vivo*. Since clonotypes were not equally represented *ex vivo*, we performed permutation tests by comparing observed distributions (the *ex vivo* barcodes of true barcodes, as detected *in vivo*) versus a null model (shuffling the *ex vivo* LARRY labels across the cells). The LSC-3_ev_ (MPP-like) clonotype emerged with the highest relative engraftment ratio (i.e. chance of any *in vivo-*detected clone to be from a particular clonotype *ex vivo*) and *in vivo* fitness (i.e. relative clonal expansion after transplantation). LSC-0_ev_ and LSC-1_ev_ (HSC-like) clonotypes also had higher than random capacity to engraft and be detectable. In contrast, LSC-2_ev_ (GMP-like) clonotypes showed diminished engraftment and expansion capacity, and LSC-4 was undetectable (**Figure 2k**). In terms of *in vivo* fitness, LSC-1_ev_ and LSC-3_ev_ showed the highest expansion rates, whereas LSC-0 produced smaller clones (**Figure 2l**). However, it was difficult to attribute this property to any specific single cluster (**Figure 2m**). To unbiasedly describe the gene programs that contribute to LSC leukemia-reinitiating potential, we grouped the clones strictly based on their *in vivo* engraftment capacity, and then performed differential expression analysis (**Figure 2n - Supplementary Table 5**). Engrafting LSCs (functional, “true”, LSCs) showed a specific gene expression program, marked by high expression of *Hoxa9*, *Rnf220*, *Cd48, H2afy*, *Tcf4*, *Hmga2*, and *Hdac9*, sharing many markers with the MPP4 subtype^62,63^. Conversely, they were depleted from markers of long-term platelet and myeloid-biased HSCs (*Vwf, Selp, Gata1, Prkg1, Hoxb9*)(**Extended Data Figure 2g**)^38,40,64–66^. At the pathway level, engrafting clones were identified by a high MAPK and PI3K axis activity, with lower trends of NF-κB and p53 pathway activation (**Figure 2o**). For a more quantitative analysis, we calculated an “engraftment probability score”, taking the embedding density of cells from engrafting clones, and subtracting the embedding density of non-engrafting clones, and then we calculated positively-correlating genes (**Extended Data Figure 2h-i**). Together, all these analyses support a model in which a diversity of stable LSC states can emerge and self-renew, but it is a subset of MPP4-like LSCs, displaying strong pro-survival signals and a reduced inflammatory tone, that harbours the strongest *in vivo* leukemia initiation capacity.

### *In vivo* variation in LSC programs is associated with stable, heritable differentiation biases

The striking, stable phenotypic variability among LSC clonotypes *ex vivo* prompted us to investigate whether similar phenomena would occur *in vivo*. We thus inspected the *in vivo* cell state heterogeneity, first using Leiden clustering and then calculating the LSC17 score to identify putative *in vivo* LSCs (**Figure 3a - Supplementary Table 6**). A small but prominent cluster (Leiden 2) was significantly enriched in LSC marker gene expression (*Cd34, Msi2, Adgrg1*), while the remaining clusters (containing the rest of the AML cells) showed different hallmarks of maturation with a large degree of myelo-monocytic diversity (GMP-like, Neutrophil-like, pDC-like, cDC-like, Monocytic-like), similar to previous reports using the *Npm1^cA^ Flt3^ITD^* model (**Figure 3b**)^27,34,67^. To compare differentiation behaviors across clones, we calculated sister-cell state similarity (**Extended Data Figure 3a**). While we observed a positive association, the overall values were low, suggesting that, in contrast to PLSTC LSC-enriched cultures, the *in vivo* AML cell manifold represents a true continuum of fate differentiation trajectories that likely emerge from LSCs towards various mature-like leukemic fates.

**Figure 3.**
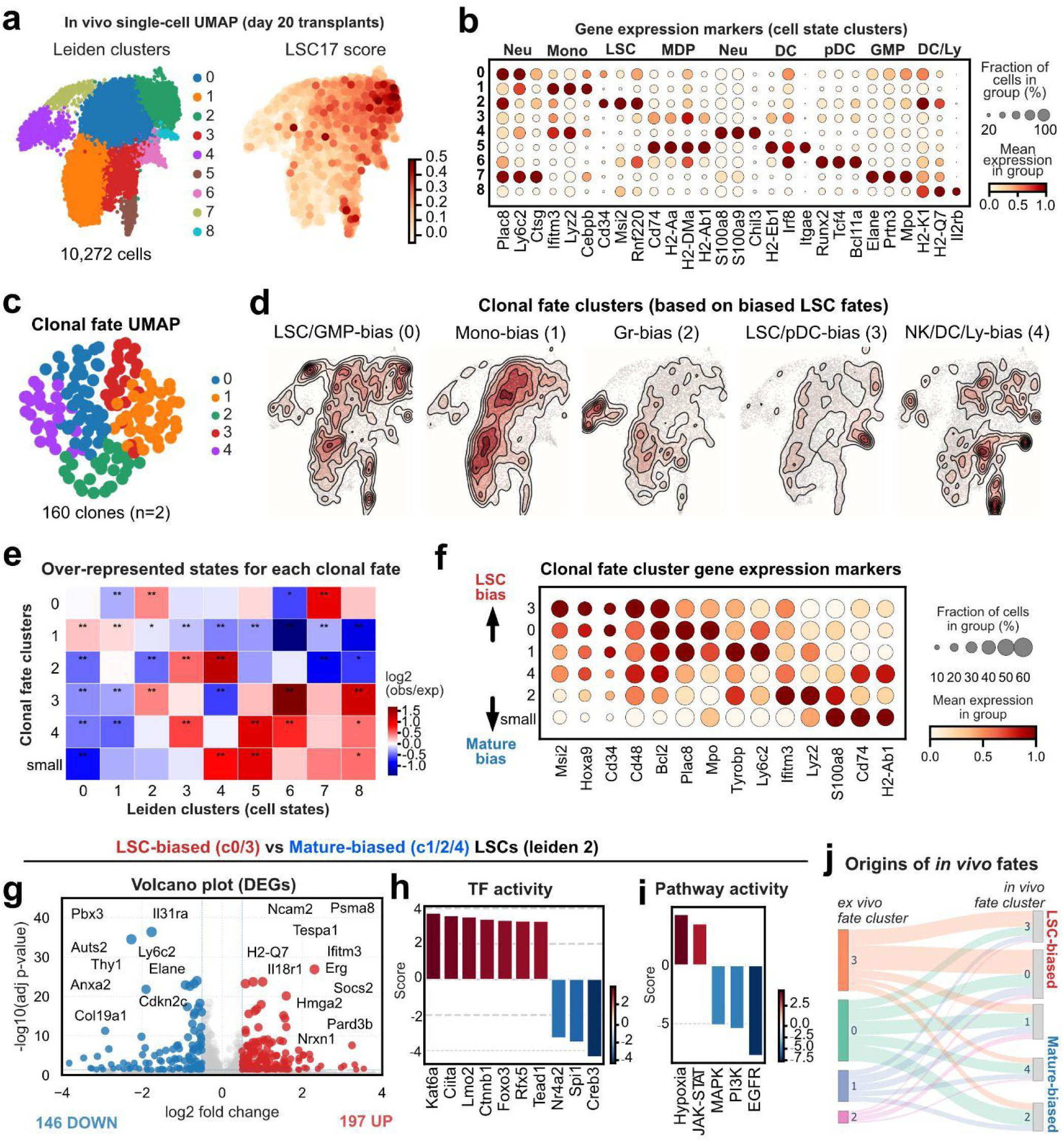
Differences in clonal LSC transcriptional programs linked to *in vivo* fate behaviors. **a**) UMAP of clonally traced *in vivo* AML cells, showing leiden cell state clusters and the calculated LSC17 gene score. **b)** Dotplot showing the *in vivo* Leiden cell state makers, and their correspondence in normal hematopoiesis. Notice the multilineage maturation that these AML cells undergo *in vivo*. **c)** Clonal fate UMAP from scLiTr, colored by fate clustering (n=160 clones from 2 mice). **d)** Density plots showing each clonal fate cluster, annotated based on their enriched states (see panel e). Notice the different biased maturation trajectories of each group. **e)** Over-representation analysis to estimate fate enrichment for every clonal cluster. Scale is log_2_ ratio of observed versus expected cell fraction in each leiden cell state cluster. Wilcoxon (multiple-test adjusted) tests, *p<0.05, **p<0.01. **f)** Clone fate cluster markers. Clusters are ordered from more LSC-biased to more mature-biased based on the proportion of cells in leiden cell state cluster 2 (LSC-like cells). **g)** Volcano plot showing differentially expressed genes within the LSC-biased versus mature-biased LSCs. Selected top markers in each group are labeled. **h)** Predicted TF activity (CollecTRI) of LSC-biased versus mature-biased LSCs, based on the results in g. **i)** Predicted pathway activity (Progeny) of LSC-biased versus mature-biased LSCs, based on the results in g. **j)** Alluvial plot connecting the clonal fate clusters *ex vivo* (left column) with their corresponding fate behavior *in vivo.* In vivo fate clusters (right column) are ordered from LSC-biased (top) to mature-biased (bottom).

To quantify the clonal heterogeneity of fate differentiation trajectories, we performed clonal fate clustering with scLiTr. Clonotype analysis grouped the clones into 5 clusters, each corresponding to a distinct group of fate-bias, suggesting that, although certain trajectories are highly conserved, different LSC clones produce unique, restricted hierarchies within the leukemic differentiation manifold, echoing the known fate heterogeneity of normal HSPCs (**Figure 3c-d**)^38,40,68–71^. Over-representation analysis identified two fate clusters (LSC-0_iv_ and LSC-3_iv_) with a distinct enrichment of the LSC state (Leiden 2), suggesting the existence of LSC-biased clones that have a diminished output of mature-like cells (**Figure 3e**). Concomitantly, the pseudo-bulk gene expression programs of clones with LSC-biased behaviors (LSC-0_iv_ and LSC-3_iv_) showed the highest expression of LSC and myeloid MPP markers (*Cd34*, *Msi2*, *Hoxa9*, *Cd48*, *Bcl2*), which were conversely reduced in the other clonotypes (**Figure 3f**). Considering that all clonotypes contain cells within the LSC state cluster (Leiden 2), we hypothesized that their different fate propensities might be explainable by gene expression differences that are already present within this most immature compartment. Differential expression analysis within the Leiden 2 cluster identified 197 differentially upregulated genes in LSC-biased (0/3_iv_) versus Mature-biased (1/2/4_iv_) LSCs (**Figure 3g and Supplementary Table 7**). The LSC-biased program showed a prominent expression of LSC genes (*Msi2, Hmga2*) as well as megakaryocytic/erythroid (MegE) progenitor markers (*Mef2c, Med12l, Rgs18, Bcl11a, Hbb-bt, Plek, Vamp5, Cd81, Coro2a, Tns1*), with high activity of JAK-STAT and Hypoxia regulatory networks (**Figure 3h-i**). This indicates unexpected parallelisms of LSCs with normal HSC heterogeneity, where platelet-biased stem cells show a strong association with self-renewal bias and slow differentiation kinetics^38,40,64,65,72,73,38–40,64,65,74–76^. We next tested whether *in vivo* LSC fate behaviors were similar across separate mice, revealing a significant degree of clonal fate stability (**Extended Data Figure 5b**). We next sought to associate different fates with pre-existing differences in their *ex vivo* LSC programs (under PLSTC conditions). While we observed a trend towards enrichment in the MPP-like LSC-3_ev_ clonotype, this was not significant compared to random label permutations (**Figure 3j**). Therefore, *in vivo* leukemic clones display stable fate heterogeneity, associated with distinct LSC gene expression patterns, and these programs are only partly determined by pre-existing differences in the *ex vivo* regulatory programs.

### Chemotherapy selects for LSC clones that engage megakaryocytic/erythroid-like programs in the spleen

Next, we asked whether the *in vivo* LSC clonal programs respond and evolve differently under the selective pressure of chemotherapy. We transplanted barcoded Npm1^cA^ Flt3^ITD^ PLSTC clones across eight sublethally-irradiated recipients (900 cGy). We confirmed leukemic phenotype by complete blood counts at day 14 post transplantation, and randomly assigned four mice to a DMSO control arm and four mice with a protocol of standard induction chemotherapy, 5 days cytarabine (araC) and 3 days doxorubicin (Doxo)(**Figure 4a**). At day 14 post-treatment (approximately 35 days post transplantation), we observed a therapeutic response, but not complete, as is frequently the case for Flt3^ITD^ leukemias. We isolated AML cells from bone marrow (BM) and spleen (SP) and prepared pooled, antibody-hashed single-cell libraries together with LARRY barcode libraries (in the BM, we pooled 50% cKit^+^ and 50% cKit^−^ sorted LARRY-T-Sapphire^+^ cells, whereas SP cells were only sorted for LARRY expression). After quality control, we detected an average of 13 clones +/− 2 per mouse (n=8 mice), with a median clone size of 19 cells. Inspection of clone sizes across samples indicated a higher Spearman correlation between treated samples compared to controls, suggesting that, while a common set of potent LSCs initiates leukemias across all mice, therapy selects a specific subset of clones (0.87 versus 0.82, two-sided t-test p=0.007, **Figure 4b, Extended Data Fig. 4a**). Single-cell transcriptome analysis revealed a continuous myeloid differentiation manifold with distinct immature (LSC-like) and mature AML cell types, which we interpreted using gene signature scores (**Figure 4c - Supplementary Table 8**). Inspecting density plots and MiloR neighborhood analyses, we found that all treated samples had robustly depleted mature progenitor states and enriched immature states (LSCs and GMP-like clusters) compared to controls (**Figure 4d-e and Extended Data Figure 4b**). This is consistent with previous findings that standard induction chemotherapy spares cells within therapy-resistant immature populations, particularly in an aggressive AML subtype such as Npm1^cA^Flt3^ITD77,78^.

**Figure 4.**
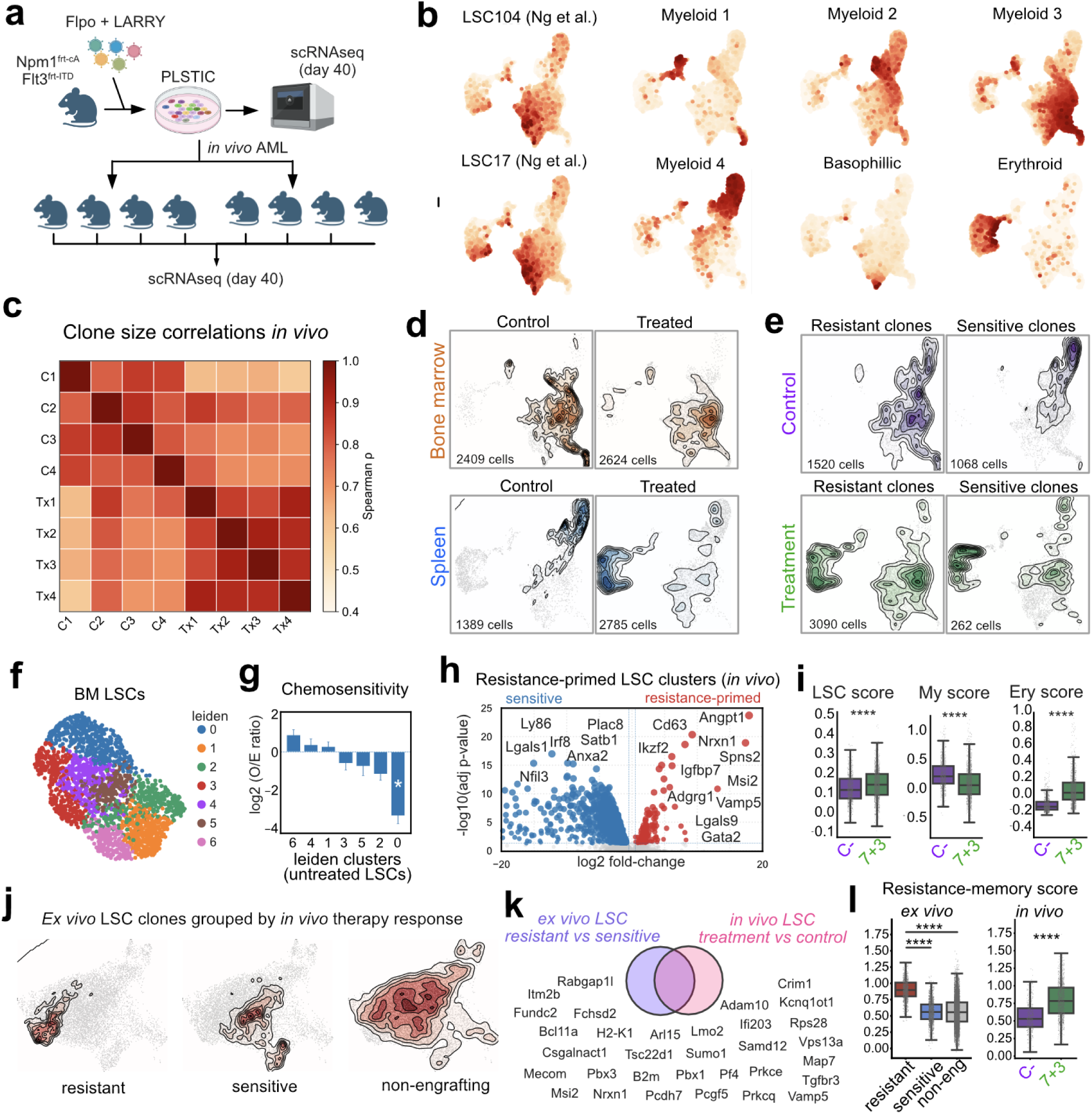
Chemotherapy selects for LSC clones that engage megakaryocytic/erythroid-like programs in the spleen. **a**) PLSTC-cultured LARRY-barcoded LSCs were expanded for 40 days and then split for profiling single-cell RNAseq or for transplanting into irradiated recipients. Recipients were monitored using tail-vein bleeding for AML engraftment. Mice with confirmation of engraftment were randomized to initiate induction chemotherapy (araC and Doxo) or DMSO control. Two days after the end of treatment, mice were euthanized, EGFP barcoded cells were sorted from spleen or bone marrow and profiled by single-cell RNAseq to capture transcriptomes and LARRY barcodes. **b)** UMAP showing expression of gene signatures scores for different mature and LSC signatures. **c)** Correlation matrix (Spearman) of LARRY clone sizes across control and treated mice. **d)** UMAP shows density plots of AML cells in the different organs (bone marrow, top, spleen, bottom) under different treatment conditions. **e)** UMAP showing density plots of AML cells from clones classified as resistant (left) or sensitive (right). **f)** Cell state landscape of LSCs subsetted from all bone marrow samples, re-clustered using Leiden. **g)** Quantification of chemosensitivity (observed vs expected ratio) of different BM LSC clusters based on random permutations using the distribution under treatment as the null hypothesis (averaged across all clones). Cluster 0 is depleted, suggesting low chemosensitivity (high chemoresistance). **h)** Differential gene expression Volcano plot comparing resistant versus sensitive LSCs (*in vivo*) in the control condition. Top selected genes are highlighted. **i)** Quantification of AML transcriptional programs using gene scores based on marker genes (comparing control and treatment (7+3) samples). ****p<0.001. **j)** UMAP of *ex vivo* PLSTC cultures sampled before transplantation (day 40 of cell culture). Density plots show distinct states for clones based on their *in vivo* behavior after transplantation and therapy: *in vivo* resistant (left), *in vivo* sensitive (middle), and non-engrafting clones (right). Clones in the left were considered “resistance-primed” **k)** Gene program overlapping between the *ex vivo* resistance-primed DEGs (top 200 genes) and the *in vivo* treated vs control DEGs (top 200 genes). The overlap (“Resistance-memory”) consists of 32 genes, which are highlighted here. **l)** Quantification of “Resistance-memory” program across *ex vivo* and *in vivo* scenarios, comparing resistant-primed and sensitive clones (*ex vivo*), and the same clones with and without 7+3 treatment (*in vivo*). ****p<0.001.

To identify how different clones were responding to therapy, we classified them into intrinsically sensitive (reduced clone size upon therapy across multiple mice) and intrinsically resistant (larger clone size upon therapy across multiple mice), and then inspected the behaviors of their LSCs (see Methods). After reclustering the BM LSCs, we discovered 7 clusters (c0-c6), which represented a continuum of LSC states across control and therapy-response (**Figure 4f, Extended Data Figure 4c-d**). Treatment-persisting LSCs were enriched in cluster 0, which is a state marked by multilineage gene expression (**Figure 4g**). LSC clones classified as “resistant” expressed a distinct signature even in the control condition, suggesting that they were intrinsically primed for therapy-resistance (**Figure 4h - Supplementary Table 9**). We noted that many of these genes included megakaryocytic-erythroid-fate markers and transcription factors, indicative of multilineage expression priming (*Lmo2*, *Gata2*, *Vamp5*). Laying over the gene signature scores for LSCs, Myeloid and Megakaryocytic-Erythroid (MegE) progenitors, we found that treated LSCs displayed increased expression of LSC signatures, and a reduced expression of Myeloid programs (**Figure 4i**). Strikingly, we also observed a very strong upregulation of MegE programs, which are unusual in NPM1-mutant AML, though mentioned in clinical pathology reports^79^. In addition to this LSC state-fate switch in the bone marrow, we also observed an unexpected enrichment of LSCs in the spleen of treated mice, which showed an even stronger enrichment for MegE gene expression (**Figure 4b-e**). Thus, we conclude that chemotherapy induces a fate-switch within LSCs, from Myeloid-to-Erythroid programs, and this correlates with an expansion of Megakaryocytic-Erythroid LSCs in the spleen.

To identify the origin of this MegE-LSC program, we used the *ex vivo* data to trace back the clones and compare the resistant and sensitive LSCs *ex vivo*. Surprisingly, resistant LSC clones showed distinct states already in the *ex vivo* cultures, suggesting a long-lived clonal program that predetermines resistance (**Figure 4j, Extended Data Figures 4e-i, and Supplementary Table 10**). Comparing the *ex vivo* resistance-primed LSCs and the treatment-induced LSC gene-expression signatures, we found a common enriched group of genes, which included certain leukemia stemness regulators (*Mecom*, *Msi2*) and a wide variety of MegEry progenitor markers (*Lmo2*, *Pf4*, *Pcdh7*, *Vamp5*, *Fundc2*) that was different from the leukemia initiation and engraftment programs (**Figure 4k, Extended Data Figure 4j-m, and Supplementary Table 11**). Surprisingly, untreated LSCs (belonging to the resistant-primed *ex vivo* clones) did not express the signature *in vivo*, yet it was significantly activated upon treatment(**Figure 4l**). This suggests a transcriptional-memory behavior, where the resistance gene program partly shuts down, becoming latent while LSCs expand and differentiate, and then it can become re-engaged after chemotherapy, possibly mediating clonal persistence.

### Targeted CROPseq identifies Csgalnact1 as a regulator of the primitive, resistant LSC program

Our clonal tracing and in vivo chemotherapy experiments suggested that a subset of LSCs is intrinsically primed to survive induction chemotherapy by prioritising a MegE-like resistance program. To decipher the genes required to maintain this LSC state, we performed a focused single-cell CROPseq screen in PLSTC cultures. We created a dual-guide RNA library using our own modified CROPseq-EF1a-T-Sapphire vector and the tRNA-architecture for dual guide expression^80,81^. We selected candidates from the various LSC signatures focusing on genes that were also expressed in human primitive AML datasets (*Csgalnact1, Adgrg1, Spns2, Zeb1, Mat2a, Zmat1*), and we added 2 essential (*Plk1, Gapdh*) and 2 non-targeting controls (NTCs). We transduced a pool of Npm1^cA^/Flt3^ITD^ PLSTCs with a constitutively expressed SFFV-SpCas9-mScarlet lentivector and the dual guide library and profiled T-Sapphire^+^/mScarlet^+^ sorted AML cells 8 days after transduction **(Figure 5a).** This design allowed us to simultaneously map the effects of each perturbation on the global LSC enriched state-space.

**Figure 5.**
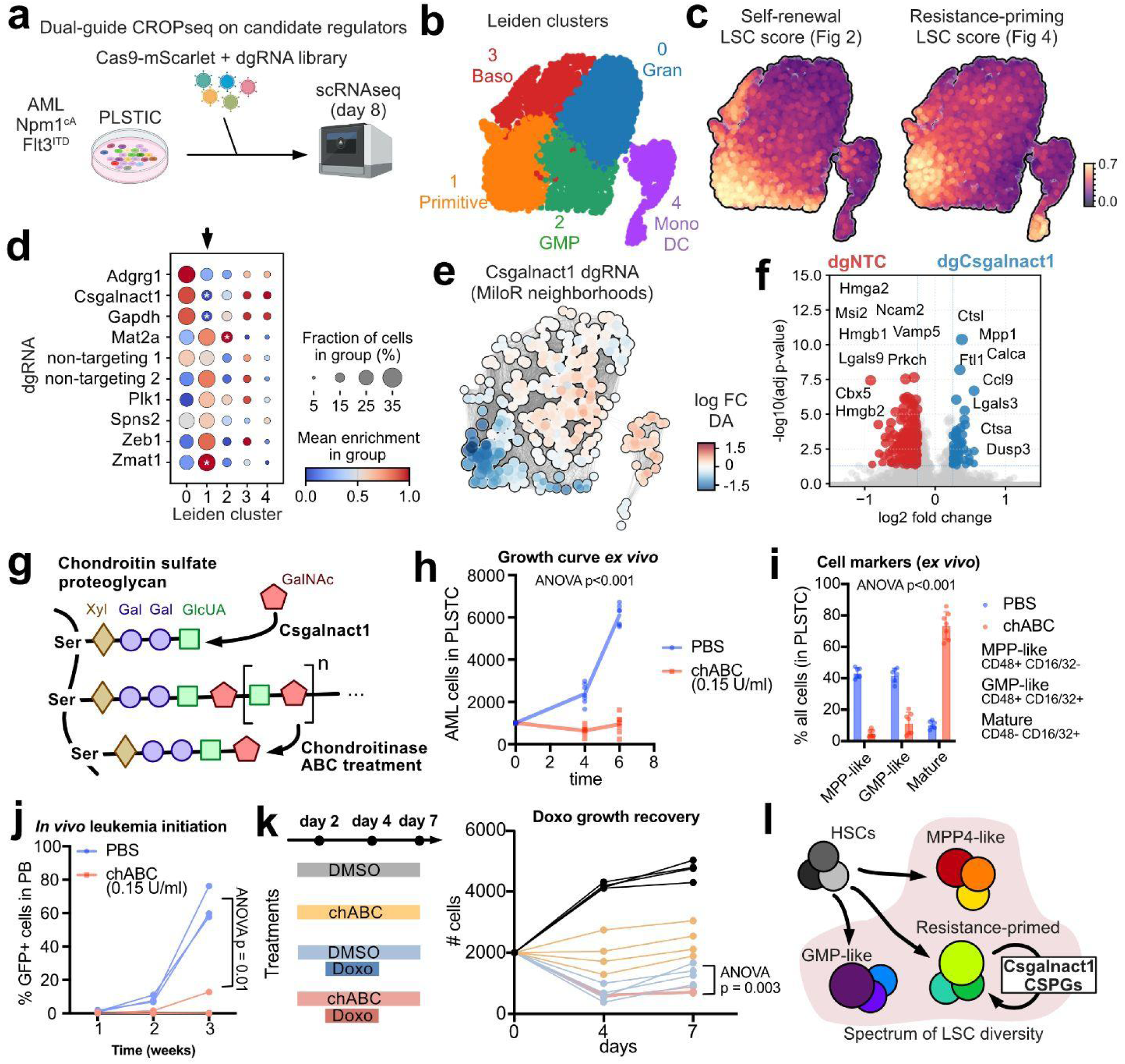
Candidate CROPseq screen in PLSTCs identifies Csgalnact1 as a regulator of the primitive-like LSC state. **a**) Experimental design. PLSTC AML LSCs were transduced with a constitutive Cas9-mScarlet lentivector and a mini dual-guide RNA library using the CROPseq architecture. At 8 days post transduction, double-positive cells were sorted and profiled by scRNAseq, sequencing transcriptomes and CROPseq guides. **b)** UMAP showing Leiden clusters of the single-cell data identified 8 programs, with 3 programs being enriched in the most immature LSC markers. **c)** UMAP showing the signature scores for the *ex vivo* resistant-primed gene signature and the *ex vivo* engrafting LSC signature. **d)** Dotplot showing the Leiden cluster (columns) enrichment score for cell groups carrying each dual-guide RNA (rows, scaled, grouped by target gene). Scale is red (enrichment) to blue (depletion). Dot size represents the fraction of cells. *p<0.05 (permutation test). **e)** Milopy UMAP, colored by neighborhood enrichment/depletion (using PCA dimensions and 100 neighbors), comparing Csgalnact1 dgRNA cells versus the rest. Scale is red (enrichment) to blue (depletion). **f)** Volcano plot showing differentially expressed genes contrasting Csgalnact1 dgRNA cells versus the rest. Selected top primitive LSC (left, downregulated) vs mature LSC (right, upregulated) genes are highlighted. **g)** Chondroitin sulfate (CS) proteoglycan synthesis pathway showing the role of Csgalnact1 and ABC Chondroitinase as a treatment. **h)** PLSTC growth curve of LSCs treated with ABC Chondroitinase (0.15 U/ml) in red and PBS control in blue (n=6 independent treatments, ANOVA p<0.001). **i)** Progenitor and stem cell marker analysis *ex vivo* in PLSTC cells treated with ABC Chondroitinase every day for 7 days (n=6 independent treatments, ANOVA p<0.001). **j)** In vivo leukemia growth capacity of control and ABC Chondroitinase-treated PLSTC cells, measured as % GFP (leukemic) cells in peripheral blood after cultured cells were transplanted into lethally irradiated mice (100 cells per mouse, n=4 per group, ANOVA p=0.01). **k)** Doxorubicin treatment response and recovery analysis of control and ABC Chondroitinase-treated PLSTC cells. n=4 independent experiments. ANOVA (Doxo vs Doxo+ABC p=0.003). **l)** Model for LSC diversity. After HSPCs acquire oncogenic mutations, each LSC clone diversifies into a distinct stable state, with unique functional properties (self-renewal, differentiation, drug resistance) and associated regulatory program. CS synthesis may be a targetable dependency of primitive, resistance-primed LSCs.

Unsupervised analysis of the CROPseq dataset recovered five Leiden clusters that recapitulated the typical PLSTC cellular distribution **(Figure 5b).** Projecting the ex vivo resistance-primed LSC signature and the self-renewal LSC signatures onto this UMAP confirmed that these two states occupy related but only partially overlapping regions **(Figure 5c).** The resistance-primed score peaked in a subset of the most primitive clusters, whereas the self-renwal score extended more broadly into both primitive and GMP-like states, consistent with our earlier observation that self-renewal and resistance properties are linked, but distinctly enriched and regulated by different programs.

We next quantified how each perturbation redistributed cells across these programs by testing the enrichment or depletion of dual-guide carrying cells within each Leiden cluster **(Figure 5d)** as well as differential abundance testing with replicates using MiloR^82^ neighborhood enrichment-analysis (**Figure 5e**). Among our targets, Adgrg1, Gapdh and Csgalnact1 stood out, being markedly depleted from the primitive LSC states. While Plk1 guides were clearly depleted as expected for a classic essential gene, Gapdh guides were not, indicating that it is not essential under the specific conditions of PLSTC media. However, glycolisis is a known metabolic dependency of HSCs and LSCs, particularly in Flt3-ITD AML^83^, which might explain the specific depletion of Gapdh guides from primitive-like LSC states. Adgrg1 encodes GPR56, a known marker of LSCs and a regulator of AML stemness, and we expected its depletion would abolish the primitive-like LSC states^51,84–86^. Finally, we found Csgalnact1 as an unexpected hit. Csgalnact1 is the main regulator of chondroitin-sulfate (CS) synthesis, catalyzing the initiation step of the sugar chain polymerization on proteoglycans. Differential abundance analysis showed that the primitive state was strongly depleted upon Csgalnact1 perturbation, which biased LSC states towards more mature granulocytic and monocytic states. Differential expression analysis indicated that Csgalnact1 disruption leads to significant downregulation of key members of the primitive and resistance-primed LSC programs (Hmga2, Msi2, Hmgb1/2, Ncam2, Vamp5, Lgals9, Cbx5, and Prkch) while inducing genes associated with partial granulomonocytic differentiation (Ctsl, Ctsa, Lgals3, Ccl9, Mpp1, Ftl1, Calca, Dusp3) (**Figure 5f - Supplementary Table 12**).

CS synthesis plays a key role in central nervous system scarring and has been the subject of intense research to treat spinal cord injury and other conditions. It has also been investigated in the context of solid tumors and even some leukemias, due to the role of the extracellular matrix in promoting tumor progression. A safe, non-toxic treatment involves the use of ABC chondroitinase, a recombinant enzyme that breaks down the CS polymer in proteoglycans (**Figure 5g**), altering their normal function. We hypothesized that ABC chondroitinase would block LSC stemness and therefore applied it to PLSTC cultures. Surprisingly, a moderate dose of ABC chondroitinase (0.2U/ml) was sufficient to completely block LSC self-renewal in PLSTC media (**Figure 5h**). We then analyzed cell surface markers in remaining non-proliferating cells, which suggested that ABC treatment forced cell differentiation (**Figure 5i**). Transplantation experiments of ABC-treated and control PLSTC cells indicate that ABC-treated cells can still engraft, although there was a trend towards slower growth (**Figure 5j**). Finally, we hypothesized that ABC-treated cells would be more sensitive to standard chemotherapy. We treated ABC and control PLSTC cells with araC and Doxo (IC80 dose, 6 nM Doxo, 6 uM araC) for 2 days, and then let them recover in normal media for an additional 3 days (**Figure 5k**). ABC-treated PLSTC cells less capable of recovering post treatment (post-Doxo growth rate of 1.17+/−0.69 for controls, compared to 0.27+/−0,26 for ABC-treated, p=0.048), suggesting that chondroitin-sulfate proteoglycans are necessary for LSC drug escape.

## Discussion

Relapse of chemoresistant cells remains the principal barrier to durable remission in acute myeloid leukemia^87^. Leukemic stem cells (LSCs), while they comprise only a minor fraction of the cancer, are central to these relapses^28,88^. A major bottleneck in developing effective treatments is the lack of experimental systems that can both expand LSCs at scale and preserve their properties and heterogeneity. Our work establishes a scalable, defined culture system that enriches and preserves functional leukemic stem cells and can be used to infer the logic behind heterogeneity and drug resistance.

In stark contrast to conventional serum-containing media that rapidly drift toward granulocytic and monocytic fates, PLSTCs maintain Npm1^cA^ AML cells in an immature LSC-enriched state, capable of sustained leukemia-initiating capacity. The key enabling reagent for PLSTCs appears to be HemEx type 9Ø media (a Soluplus-containing F12 nutrient mix media), since we have not been able to attain similar results with the earlier versions of Polyvinyl Alcohol (PVA)-F12 media commonly used for HSC expansion. The other key enabling aspect is to maintain the LSCs in 96-well format, and we have not been able to expand them efficiently in larger formats, possibly due to key cell-cell signaling that requires a minimum volume and cell density. Once established, the cultures are amenable to freezing and thawing multiple times, and we have been able to expand LSCs up to 10,000-fold through more than 10 serial splittings, without alterations in phenotypic states. Moreover, we have preliminary evidence that electroporation and puromycin-selection is also possible in this model. It is clear that PLSTCs would unlock complex and advanced genetic engineering (such as gene knock-ins) circumventing the need to generate new mouse models for every research question. We anticipate PLSTCs will emulate the recent success in pancreatic cancer research thanks to shareable primary adenocarcinoma cell lines^89^. As expected, PLSTC-expanded cells show reduced sensitivity to cell-cycle dependent chemotherapy and heightened sensitivity to metabolic therapy, suggesting it is an excellent primary-cell model of primitive AML states, which are challenging to model outside the *in vivo* context^10^. In the future, this could help create specific high-throughput screens that circumvent the need for precious patient material and avoid the modeling limitations of established AML cell lines. While so far we have developed PLSTC cultures only for mouse cells, it is foreseeable that similar approaches will result in successful LSC-enriched human culture models.

We have also generated new, multi-indexed EGFP LARRY barcoding libraries, which enabled us to multiplex donors and study intrinsic, heritable LSC programs across hundreds of clones, which differed in their functional properties, from *ex vivo* expansion to *in vivo* therapy resistance. This functional diversity evokes the variability of patient LSCs in NSG transplantation models^90^. LSC clonotypes appear to display slow fluctuations, causing the same clone to remain confined within restricted regions of the transcriptional manifold across weeks of culture and even transplantation. This clonal transcriptomic stability evokes the different trajectories during cell reprogramming^91^, as well as the emergence of multiple stable cancer cell states upon drug treatment^92^. Relative stability of heterogeneous states is a prerequisite for selection of non-genetic adaptations to stress, a phenomenon that likely underpins cancer evolution in response to therapy^93–95^. In this study, AML states showed higher stability than those that we recently reported in the pre-leukemic setting^16^, We speculate that Flt3^ITD^ mutations unlock an increase in phenotypic stability, which may underpin the therapy-resistance of leukemias carrying this driver mutation^96^. Further work will be needed to establish the precise signaling and mechanisms downstream of Flt3^ITD^ that drive increased phenotypic stability.

Tracking clones through chemotherapy in vivo, we found that surviving LSCs accumulate in a specific, rare cell state, marked by megakaryocytic-erythroid (MegE) progenitor markers, while being depleted from myeloid states. Surprisingly, resistant clones already occupy a distinct transcriptomic state *ex vivo*, weeks before the cells are injected to generate leukemias and treatment is performed. This LSC program is lost as clones reinitiate leukemia during engraftment *in vivo*. But then, cells appear to re-enter this state during therapy, which is accompanied by expansion of a novel MegE AML trajectory in the spleen. These results suggest a therapy-specific phenotypic switch, with cells acquiring a primitive-like, multipotent myelo-erythroid state that is permissive in our *ex vivo* conditions but rare during normal leukemogenesis. Alternatively, this switch may be driven by the selection of pre-existing, ultra-rare, resistance-primed LSCs (based on gene score, 1/50th of all LSCs in absence of therapy), and then they become massively selected upon treatment. However, considering their initial rarity and their high abundance post-therapy, these LSCs would have to cycle ∼2 times per day (while under treatment), to reach the numbers that we observe in our experiment. We consider these cycling rates unlikely, particularly in light of their mostly quiescent-like state, and, therefore, our current evidence seems to favor the transcriptional-memory hypothesis. We posit that certain LSC clones can retain hidden fate potentials, and these can be released upon stress, to survive therapeutic challenges. In the future, temporal single-cell profiling of LSC lineage, DNA methylation, chromatin, and transcriptomic states will uncover the layers of these latent resistant programs.

In our study, we prioritized uncovering therapeutically-available targets with expression conserved in human LSCs. We leveraged a combination of PLSTCs and a new dual-guide CROPseq strategy (similar to^80^, but of our own design), and we unveiled the unexpected role of chondroitin sulfate synthesis. Csgalnact1 is the main enzyme that drives initiation of chondroitin sulfate chains on proteoglycan cores^97,98^. Proteoglycans regulate a variety of cell and tissue properties, shaping cell–cell and cell–matrix interactions, growth factor availability, and niche signaling^99^, and their remarkable diversity unravels in hematopoiesis^100,101^. Proteoglycans can be modified through different glycosaminoglycans (GAGs). Two GAGs, Heparan sulfate (HS) and Chondroitin sulfate (CS) share the same initiation pathway, with the rate of CS elongation being determined by Csgalnact1 expression. Our working model is that LSCs leverage Csgalnact1 to change the CS/HS balance in key proteoglycans facilitating growth factor-independent or autocrine signaling to sustain the self-renewal of the primitive-like state. Supporting this view, functional overexpression of CSGALNACT1 confers cytokine-independent growth in Ba/F3 cell line models^102^. A recent paper described the role of CS proteoglycan CSPG4 (also called NG2) in MLL-rearranged Acute Lymphoblastic Leukemias^103^, but we could not detect CSPG4 expression in our NPM1 AML. Therefore, the specific proteoglycans that are involved in the maintenance of the primitive AML LSC state (downstream of CSGALNACT1) remain a top priority for future research efforts.

Could CS synthesis thus be a new target for treating AML? CSGALNACT1 is expressed in normal mouse and human hematopoiesis and is particularly enriched in HSCs and MPPs. Csgalnact1 knock-out mice show a mild phenotype at steady state, likely due to its compensation by Csgalnact2, suggesting it could be a safe target^99,104^. Loss of Csgalnact1 impairs long-term engraftment capacity, but only upon serial transplantation, indicating its role in regeneration under stress^105^. Our results are consistent with this, suggesting that CS perturbation selectively depletes the highly self-renewing, chemoresistant LSCs, while sparing more committed monocytic/granulocytic AML programs. Elevated CSGALNACT1 expression has been associated with high-risk disease and adverse outcomes in human lymphoma and AML. Compared to other AML subtypes, CSGALNACT1 appears to be downregulated in NPM1-mutant AML when analyzed at the bulk level^106^, and this is consistent with the dominant mature myelomonocytic compartment that characterizes these leukemias. In contrast, our inspection of publicly available single-cell RNAseq atlases suggests that CSGALNACT1, along with other established LSC markers, is distinctly expressed within a rare, primitive, stem-like state^14^, which would be consistent with our observations *in vivo*. We posit that the conserved expression pattern of CSGALNACT1 will translate into conserved mechanistic regulation, but confirmation will require specialized translational efforts.

Looking forward, we expect that PLSTC cultures will continue to unlock mutant stem cell expansion in a variety of additional leukemic conditions. In combination with advanced genome-engineering tools, we envision a future in which we can partially bypass the generation of new mouse models, and directly engineer primary LSC cultures to trace and perturb AML *ex vivo* and *in vivo*, significantly speeding up research for these deadly disorders. Finally, in a more conceptual note, our results highlight the potential of combined lineage and single-cell analysis towards a mechanistic understanding of how long-lived, stable phenotypic variation arises and contributes to disease progression and therapy response.

## Limitations

Our experiments focus on a single, genetically defined Npm1cA/Flt3ITD model; whether analogous resistance-primed programs generalize to other AML genotypes and to human primary LSCs remains to be tested. Second, our chemotherapy regimen models induction therapy but not targeted agents such as venetoclax or menin inhibitors, which may impose other selective pressures on the LSC program space. Finally, PLSTCs focus on the expansion and enrichment of primitive-like states, which are not always the main substrate of therapeutic selection (e.g. monocytic-relapses in response to Venetoclax treatment). We anticipate that a combination of culture systems will be required to comprehensively model different aspects of disease evolution and identify new targets to treat or avoid drug resistance.

## EXTENDED DATA FIGURES

**Extended Data Figure 1.**
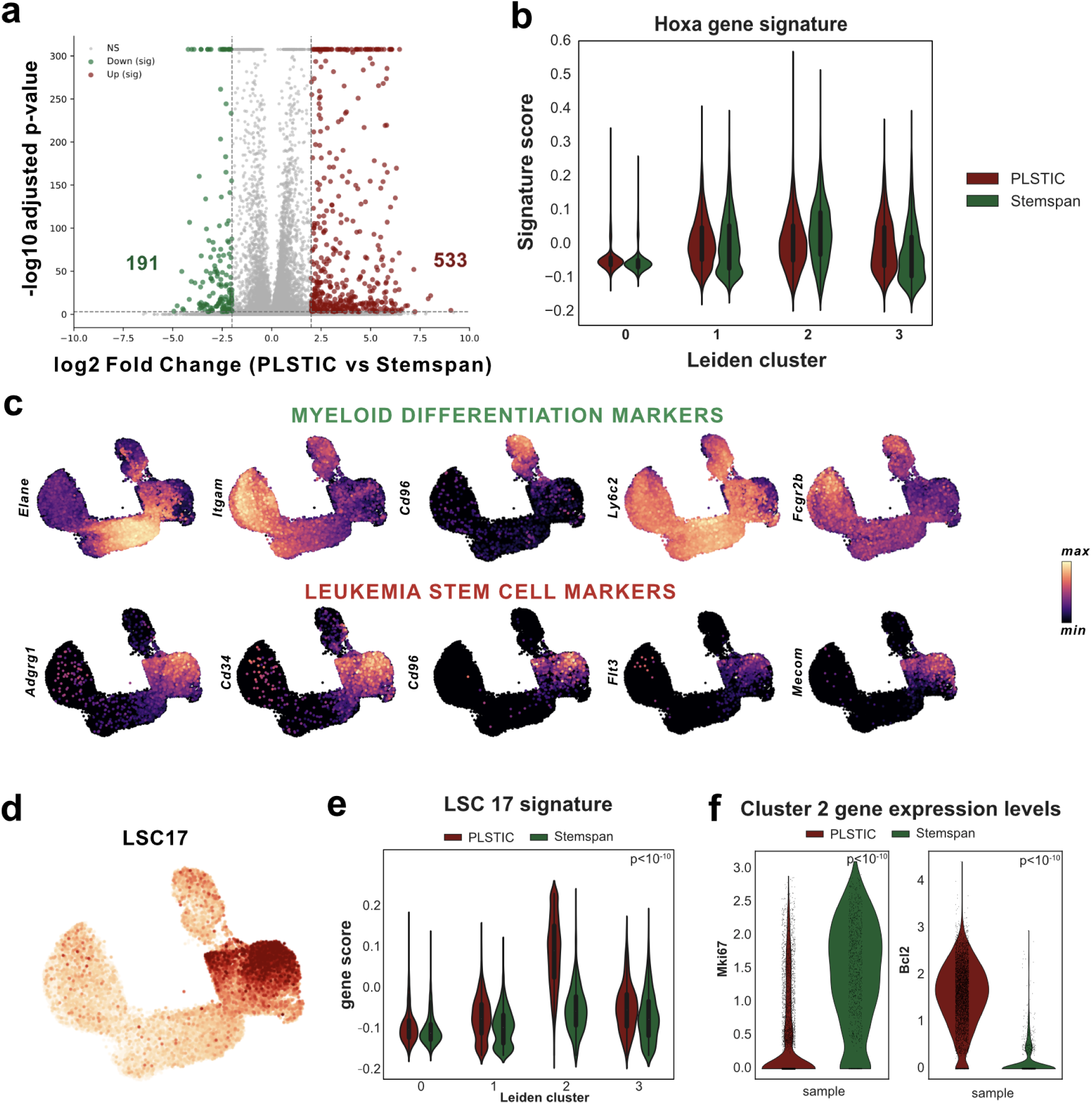
Additional analyses on PLSTC vs Stemspan cultures. **a**) Volcano plot showing differentially expressed genes in cluster 2 comparing Stemspan and PLSTC cultures. Genes with more than 2 fold difference are highlighted, indicating a substantial difference in this primitive-like cluster between the two conditions. **b)** Violin plot comparing the Hox-signature score across both conditions in all clusters, suggesting that both conditions rely on Hox-program mediated immortalization for sustained expansion. **c)** UMAP plots showing selected LSC and AML markers. **d)** UMAP plots showing the complementary LSC17 signature score. **e)** Violin plot showing the LSC17 signature score across all clusters for each condition. Wilcoxon test p-values for cluster 2 are shown. **f)** Violin plots showing the reduced expression of cycling marker *Mki67* and Venetoclax-target *Bcl2* in PLSTC versus Stemspan condition. Wilcoxon test p-values are shown.

**Extended Data Figure 2.**
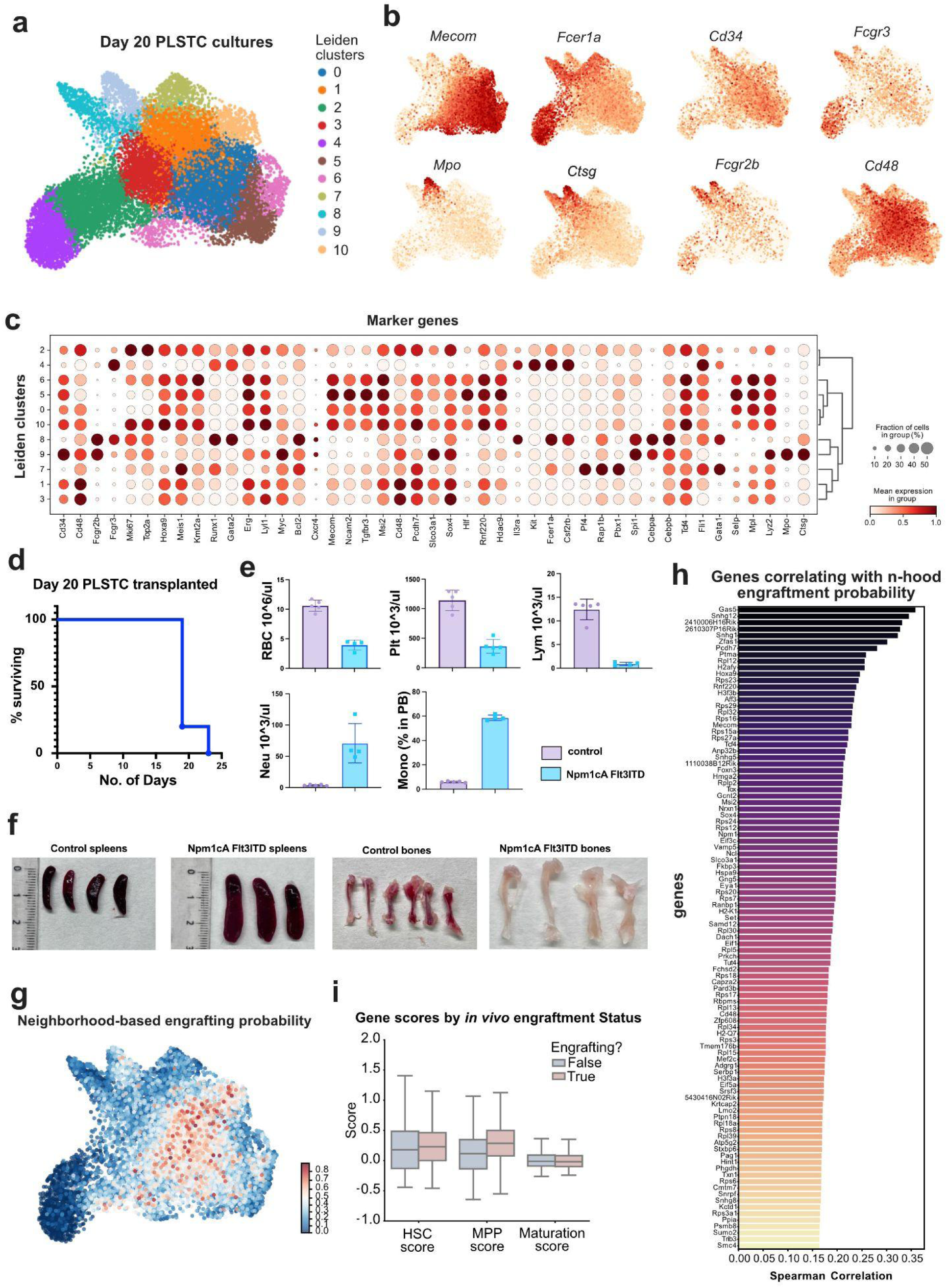
Additional analyses on day 20 PLSTC LARRY barcoding experiment. **a**) UMAP showing Leiden clustering of day 20 barcoded PLSTC leukemic cell states. **b)** UMAPs showing selected gene expression distribution (min to max scale) **c)** Dotplot showing expression levels for selected genes across different Leiden clusters **d)** Survival curves of sublethally-irradiated mice transplanted with day 20 PLSTC cells **e)** Complete blood counts analysis of peripheral blood at day 18 post transplantation. **f)** Images of representative spleens and bone marrows of control (wild-type CD45.1) and PLSTC-transplanted mice. **g)** UMAP showing the neighborhood-based calculation of engraftment probability score (neighbor-averaged embedding density for cells belonging to engrafting clones minus embedding density of non-engrafting clones). **h)** Top genes with positive cell-to-cell correlation with the engraftment probability score (Spearman). **i)** Gene scores calculated for HSC, MPP and Mature myeloid gene sets, separated by engraftment status (i.e. clone detected after transplantation).

**Extended Data Figure 3.**
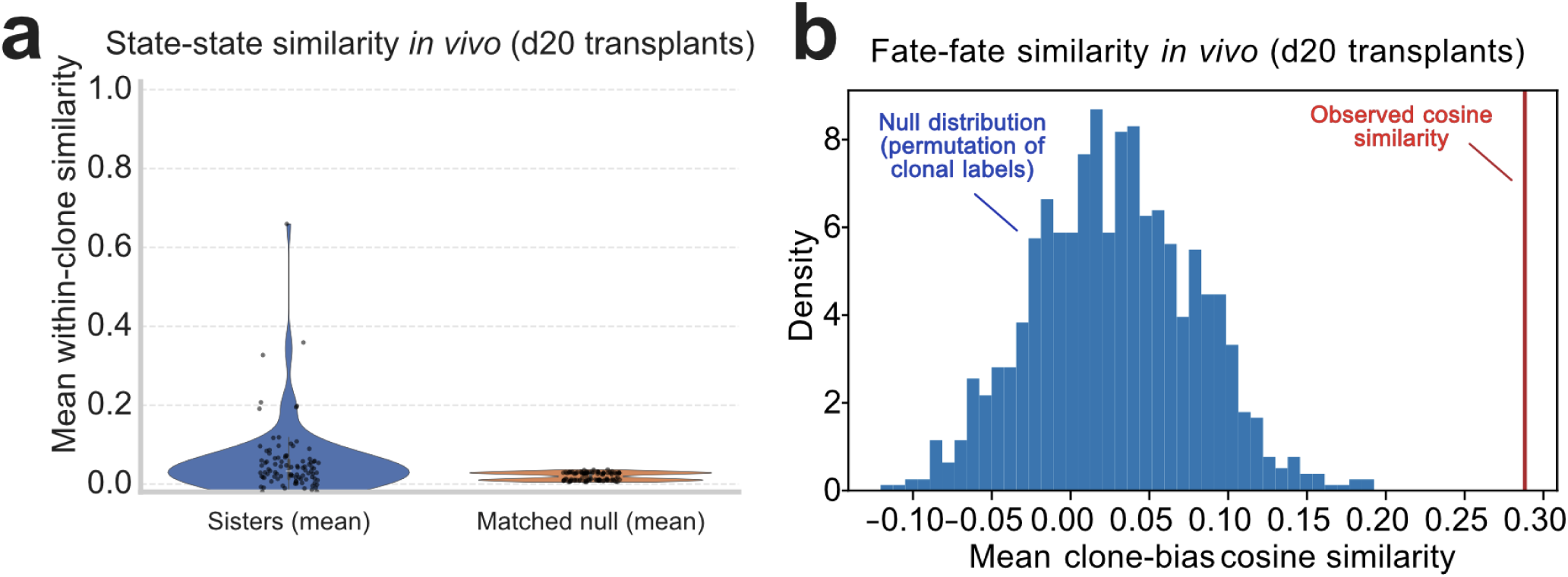
Additional data analyses on *in vivo* day 20-transplanted PLSTC LARRY cells. **a**) Sister cell cosine similarity comparing true sisters (left) versus matched label-permuted sisters (right). **b)** Mouse-mouse cosine similarity of leukemic cell fates (fate-matrix was calculated first by normalizing frequencies for each leiden cluster, separately for each mouse, and then by scaling row frequencies for every clone in each mouse).

**Extended Data Figure 4.**
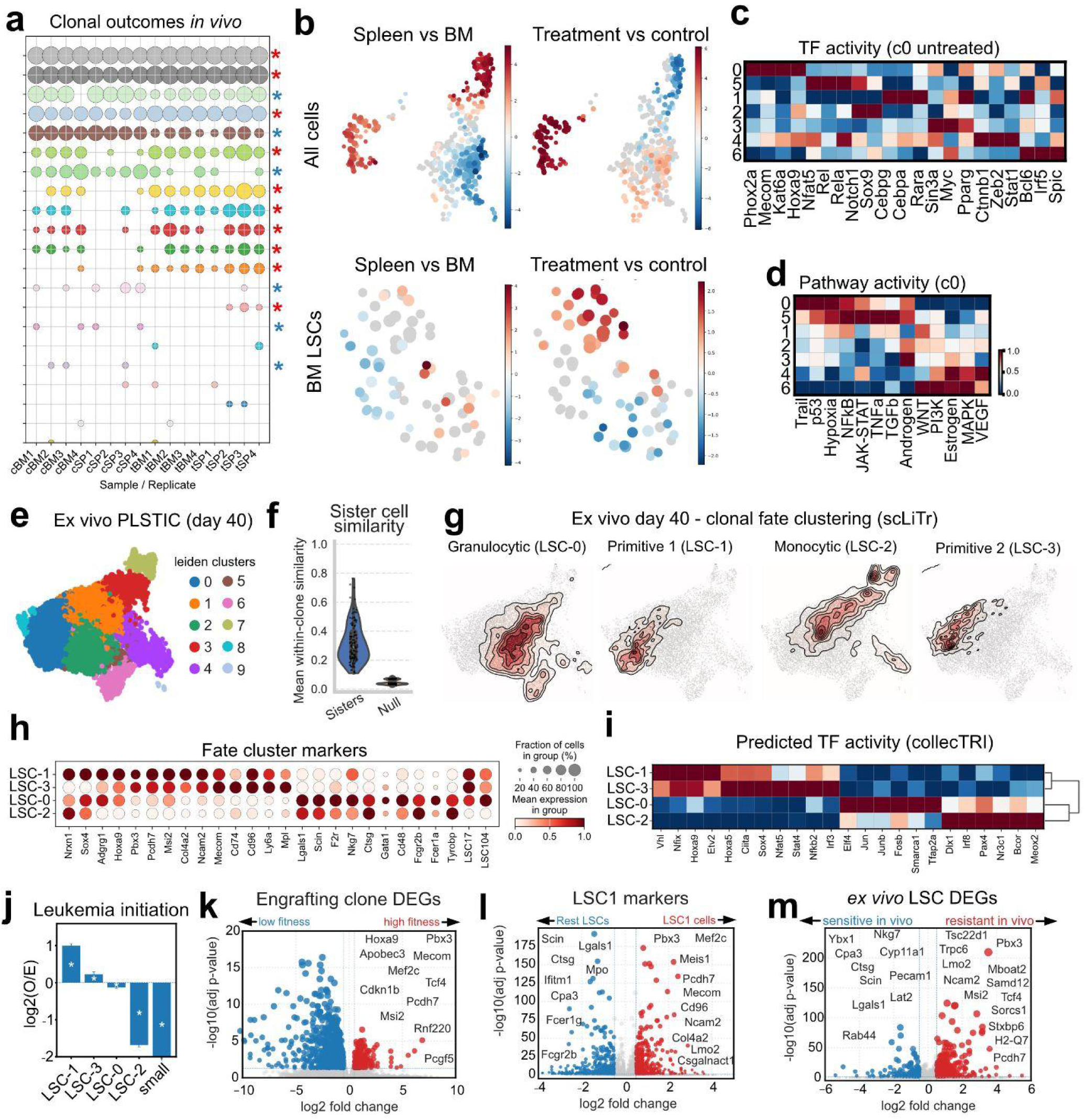
Additional data analyses on *in vivo* control and treated PLSTC-derived AML. **a**) Bubble plot showing the LARRY clone size distribution of leukemic cells across different recipient mice. Red asterisks indicate clones that increase in size upon treatment. Blue asterisks indicate clones that decrease in size upon treatment. **b)** UMAP showing the MiloR neighborhood analysis for sample enrichment, leveraging the different mouse samples (hashed using TotalSeq) for replicates. Scale is from red (enrichment in the indicated sample class) to blue. **c)** Heatmap showing the predicted CollecTRI transcription factor activity score (based on Decoupler, univariate linear models) across different LSC state clusters. **d)** Heatmap showing the predicted Progeny pathway activity score (based on Decoupler, univariate linear models) across different LSC state clusters. **e)** UMAP of *ex vivo* PLSTC cell states (day 40). **f)** Sister cell cosine similarity for all clones in PLSTC cultures at day 40, before transplantation. **g)** UMAP showing the embedding density for the different fate clusters identified by scLiTr (day 40 clones). **h)** Dotplot showing differentially expressed genes across the scLiTr LSC clonotypes (day 40). **i)** Heatmap showing the predicted CollecTRI transcription factor activity score (based on Decoupler, univariate linear models) across different *ex vivo* LSC clonotypes (day 40). **j)** Leukemia initiation potential of different LSC clonotypes at day 40 (calculated as observed versus expected ratio using clonotype label permutations to create a null model). **k)** Volcano plot showing differentially expressed genes for day 40 engrafting clones (comparing high fitness - detectable, and low fitness - undetectable clones). Cells belonging to clones detectable in any *in vivo* mouse sample after transplantation are in red (right). All other cells are in blue (left). **l)** Volcano plot showing differentially expressed genes in *ex vivo* LSC1 clonotype cells (day 40). Cells belonging to clones classified as LSC1 by scLiTr are in red (right). Cells belonging to clones not classified as LSC1 are in blue (left). LSC1 with all the other cells. **m)** Volcano plot showing differentially expressed genes within *ex vivo* LSCs (day 40) according to their therapeutic response *in vivo* after engraftment. Cells belonging to clones that engraft and resist therapy *in vivo* are in red (right). Cells belonging to clones that engraft but diminish upon therapy *in vivo* are in blue (left).

## Data and Code availability statement

Preprocessed data files and Jupyter notebooks are available at Zenodo (doi.org/10.5281/zenodo.18444428). All the raw and processed files are available through Gene Expression Omnibus (GSE318348).

## Acknowledgments

This work was supported by the Cris Cancer Foundation Excellence Award (PR_EX_2020-24), the European Commission’s Horizon Europe program, through the ERC Starting Grant MemOriStem (101042992), the Spanish National Research Agency (PID2020-114638RA-I00, PID2023-148073OB-I00), the Agencia de Gestio d’Ajuts Universitaris i de Recerca (2021 SGR 01586), and the CERCA Program/Generalitat de Catalunya. A.E.R.-F. acknowledges support from the Institut Catalá de Recerca i Estudis Avançats (ICREA), the EMBO Young Investigator Programme, and past support from the Ministry of Science Ramon y Cajal Fellowship and the LaCaixa Junior Fellows Incoming Fellowship. I.S. was supported through the European Union’s Horizon 2020 research and innovation program under the Marie Skłodowska-Curie grant agreement no 945352. The authors would like to acknowledge the technical assistance of the Functional Genomics Core Facility at IRB Barcelona (single-cell library preparations and sequencing), the Flow Cytometry and Cell Sorting Facility at the University of Barcelona (CCIT-UB), and other facilities from IRB Barcelona and Parc Cientific de Barcelona (PCB). The authors also wish to thank colleagues in the ISEH community, the Fraticelli lab, and IRB Barcelona for various helpful discussions. Illustrations were created with Biorender.

## Author contributions

A.R.F., R.B. and I.S. conceptualized the project. I.S. performed molecular biology, library preparations, cell isolation, cell culture, and animal experiments for results in Figures 2, 3 and 4. A.P. and A.M.P. performed molecular biology, library preparations, cell isolation, cell culture, and animal experiments for results in Figures 1 and 5. A.R.F. and I.S. generated the analytic pipelines and performed bioinformatic, single-cell sequencing, and statistical analyses. P.S.S. and D.F.P. assisted with data pre-processing and data analysis. I.S., M.L.O. and C.B. generated the LARRY vectors and produced the libraries. M.G., L.G. and C.G. generated and validated the CROPseq dual-guide vectors and libraries. R.B. generated the mouse crosses, provided the transgenic mouse bone marrow samples, provided guidelines for leukemia experiments, and analyzed results. A.R.F., I.S., and A.P. wrote the manuscript. All authors provided feedback and input to finalize the manuscript text. A.R.F. supervised the project.

## Material and Methods

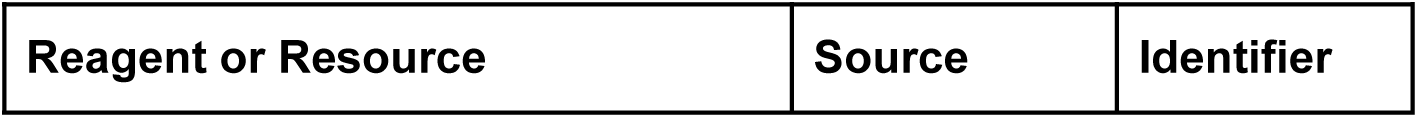

## Antibodies

|  |  |  |
| --- | --- | --- |
| Anti-CD117 Microbeads mouse | Miltenyi Biotec | 130-091-224 |
| PE anti-mouse CD201 | BioLegend | 141504 |
| APC anti-mouse c-Kit | BioLegend | 135108 |
| PE Cy7 anti-mouse Ly-6A/E (Sca-1) | BioLegend | 108114 |
| APC Cy7 anti-mouse CD48 | BioLegend | 103432 |
| PE Cy5 anti-mouse CD150 | BioLegend | 115912 |
| Pacific Blue Lineage Cocktail | BioLegend | 133310 |
| PE/Cy7 anti-mouse CD34 | BioLegend | 128618 |
| APC/Cy7 anti-mouse CD16/32 | BioLegend | 101328 |
| TotalSeq B0301 anti-mouse Hashtag1 | BioLegend | 155831 |
| TotalSeq B0301 anti-mouse Hashtag2 | BioLegend | 155833 |
| TotalSeq B0301 anti-mouse Hashtag3 | BioLegend | 155835 |
| TotalSeq B0301 anti-mouse Hashtag4 | BioLegend | 155837 |
| TotalSeq B0301 anti-mouse Hashtag5 | BioLegend | 155839 |
| TotalSeq B0301 anti-mouse Hashtag6 | BioLegend | 155841 |
| TotalSeq B0301 anti-mouse Hashtag7 | BioLegend | 155843 |
| TotalSeq B0301 anti-mouse Hashtag8 | BioLegend | 155845 |
| TotalSeq B0301 anti-mouse Hashtag9 | BioLegend | 155847 |
| TotalSeq B0301 anti-mouse Hashtag10 | BioLegend | 155849 |

## Chemicals, peptides, and recombinant proteins

|  |  |  |
| --- | --- | --- |
| HEPES buffer solution | Gibco | 15630-056 |
| Insulin-Transferrin-Selenium-X | Gibco | 51500-056 |
| Poly (vinyl alcohol) | Sigma Aldrich | 363081-1Kg |
| F-12 Nutrient Mix (Ham) | Gibco | 21765-029 |
| Direct PCR tail | Viagen | 102-T |
| IBIAN-Proteinase K solution | IBIAN lab | 106-PK01 |
| LE Agarose | SeaKem | 50004 |
| LB Agar | PanReac<br>AppliChem | A0927,1000K<br>G |
| LB Medium | PanReac<br>AppliChem | A0954,1000K<br>G |
| NEB Builder Hifi DNA Assembly<br>master Mix | New England<br>Biolabs | M5520AA |
| KAPA2G Robust hotstar Ready Mix | KAPA<br>Biosystems | KK5702 |
| KAPA HiFi Hotstart Ready Mix | Roche | 7958935001 |
| Carbenicillin Disodium Salt | Thermo<br>Scientific | 10177-012 |
| AMPure XP beads | Beckman<br>Coulter | A63881 |
| Fibronectin | R&D System | 1030-FN-05M |
| Recombinant Murine TPO | Peprotech | 315-14 |
| Recombinant Murine SCF | Peprotech | 250-03 |
| Recombinant Murine IL-3 | Peprotech | 213-13 |
| Recombinant Murine IL-6 | Peprotech | 216-16 |
| Tamoxifen,4-Hydroxy-(Z) 5 mg | EMD Millipore | 579002 |
| Tamoxifen 5G | Sigma | T5648 |
| Gibson Assembly® Master Mix | New England<br>Biolabs | E2611L |
| NEB® 10-beta Electrocompetent E.<br>coli | New England<br>Biolabs | C3020K |
| EcoRV-HF® | New England<br>Biolabs | R3195L |

## Critical commercial assays

|  |  |  |
| --- | --- | --- |
| Chromium Next GEM Single Cell 3'<br>Kit v3.1 | 10×<br>Genomics | 1000268 |
| Chromium Next GEM Chip G Single<br>Cell Kit | 10×<br>Genomics | 1000127 |
| Dual Index Kit TT Set A | 10×<br>Genomics | 1000215 |
| Dual Index Kit NT Set A | 10×<br>Genomics | 1000250 |
| NucleoBond Xtra Midi | Macherey-Na<br>gel | 740410.1 |
| NucleoBond Xtra Maxi | Macherey-Na<br>gel | 740414.5 |
| Nucleospin Plasmid | Macherey-Na<br>gel | 740588.25 |
| Monarch Genomic DNA purification<br>kit | New England<br>Biolabs | 10104305 |

## Mice and Animal Guidelines

All procedures involving animals adhered to the pertinent regulations and guidelines. Approval and oversight for all protocols and strains of mice were granted by the Institutional Review Board and the Institutional Animal Care and Use Committee at Parc Científic de Barcelona under protocol CEEA-PCB-22-001-ARF. The study follows all relevant regulations from European and Spanish law. Mice were kept under specific pathogen-free conditions for all experiments. Male C57BL6/J mice (wild-type) and Ptprca (CD45.1) mice were obtained through Charles River Laboratories, France, and conditioned using sub-lethal X-ray irradiation (900 cGy, split dose) in the morning before transplantation in the afternoon of the same day. Transplanted bone marrow cells from leukemic mouse models were from frozen vials, as explained below. All alleles (*Rosa26-Flpo^ERT2107^*, Npm1-frt-cA^108^, Flt3-frt-ITD^27^, *Rosa26-FLTG*^109^) are available through The Jackson Laboratory (strain numbers: 019016, 033164, 036725, 026932).

## PLSTC versus StemSpan experiments

We transplanted lethally-irradiated CD45.1 mice (11 Gy total, 5.5 Gy split dose) with *Rosa26-Flpo^ERT2^*, *Npm1^frt-cA^*, *Flt3*^frt−ITD^, *Rosa26-FLTG* whole bone marrow cells (10^6^ cells per recipient) and waited for full bone marrow reconstitution (16 weeks). Reconstituted mice were then injected with 75 mg tamoxifen/kg body weight by intraperitoneal injection every day for a total of 3 doses post 2 months of transplant and thereafter monitored by % of TdTomato cells in the periphery for disease progression^27^. Mice with more than 80% of Td tomato cells were considered to have growing leukemic cells and were euthanized, and bone marrow (femur, tibia, and pelvis) and spleen cells were isolated by mechanical dissociation. Cell suspensions were filtered through a 100-μm strainer, washed in ice-cold EasySep buffer (PBS + 2% FBS), and subjected to red blood cell lysis (BioLegend, Cat. no. 420302). Cells were blocked with mouse FcX-block (Biolegend) and then stained with PE/Cy5 anti-mouse FLT3 (BioLegend, 135312), APC anti-mouse CD96 (131711), PE/Cy7 anti-mouse CD34 (128618), Pacific Blue anti-mouse FcεRIα (134313), and APC/Cy7 anti-mouse CD16/32 (101328). Fluorescence-activated cell sorting (FACS) was performed to isolate cells with low or absent CD16/32 and FcεRIα expression to enrich for primitive-like LSCs^110^. Sorted leukemic cells were subsequently cultured as described below. After washing, ∼20,000 cells were transferred into flat-bottom 96-well plate wells with 200 μl of complete PLSTC media (HemEx type 9Ø media, IWAI North America Inc., cat. number A5P10P01C, supplemented with 100 ng/ml mTPO, 50 ng/ml mSCF, 10 ng/ml mIL-3, and 10 ng/ml mIL-6) or conventional serum free media (StemSpan, Stemcell technologies Cat. 09655, supplemented with the same cytokines for comparison). Cells were grown at 37°C with 5% CO2 and split 1 to 4 before reaching confluence (usually at 60-70% confluence) every 4 days for up to 90 days. For growth curve analyses, viable cells were counted every 48 hours using Trypan Blue (Gibco, 15250061) and a Countess 3 Automated Cell Counter (Thermo Fisher Scientific). Viable cell number was then divided by the number of plated cells, to calculate the normalized growth. For drug resistance assays, cells were plated in quadruplicate and treated 24 hours later with Doxorubicin hydrochloride (Sigma-Aldrich, D1515-10MG), Cytarabine (Sigma-Aldrich, C3350000), or Venetoclax (Enamine, 1257044-40-8) at the indicated concentrations. Cell viability was assessed 48 hours post-treatment by DAPI staining and analyzed on an Aurora Spectral Flow Cytometer (Cytek Biosciences).

## Leukemic stem cell preservation and cold-storage

Freshly isolated cells from bone marrow or FACS, or LSCs in cell culture were resuspended, pelleted using 15 ml centrifuge tubes (800g for 7 minutes) and resuspended in ice-cold freezing medium (90% FBS, 10% DMSO), aliquoted in 100,000-cell aliquots (100 µl each). Cryotubes were immediately flash-frozen on dry ice before being transferred directly to −80 °C for 24h and subsequently moved to liquid nitrogen tanks for long-term storage. For thawing tubes, this was done quickly to avoid DMSO-related toxicity, resuspending the cells in 10 ml PBS 2% FBS, centrifuging, and resuspending into 1 ml PLSTC media to seed 5 wells (approximately 20,000 cells per well). Media was changed the day after, to further remove DMSO traces.

## Hematopoietic stem cell isolation

Frozen vials containing 10^7^ *Rosa26-Flpo^ERT2^*, *Npm1^frt-cA^ Flt3*^frt−ITD^ whole bone marrow mononuclear cells (from uninduced mice) were thawed at 37 degrees, washed, and then sieved through a 100-μm strainer and cleansed with a cold ‘Easy Sep’ buffer containing PBS with 2% fetal bovine serum (FBS). Mature lineage cells were selectively depleted through the Lineage Cell Depletion Kit, mouse (Miltenyi Biotec, Catalog no. 130-110-470), while the resulting Lin- (lineage-negative) fraction was then enriched for c-Kit expression using CD117 MicroBeads (Miltenyi Biotec, Catalog no 130-091-224). These cKit-enriched cells were washed, blocked with FcX and stained with following fluorescently labeled antibodies: APC anti-mouse CD117, clone ACK2 (Biolegend catalog no. 105812), PE/Cy7 anti-mouse Ly6a (Sca-1) (Biolegend, catalog no. 108114); Pacific Blue anti-mouse Lineage Cocktail (Biolegend, catalog no. 133310); PE anti-mouse CD201 (EPCR) (Biolegend, catalog no. 141504); PE/Cy5 anti-mouse CD150 (SLAM) (Biolegend, catalog no. 115912); APC/Cyanine7 anti-mouse CD48 (Biolegend, catalog no. 103432). EPCR^+^Lineage^−^Sca-1^+^cKit^+^CD48^−^CD150^+^ bone marrow cells were sorted by FACS with a BD FACSAria Fusion with a 100 µm nozzle and seeded into 96-well plates.

## Polymer-based cell cultures

*Ex-vivo* cultures of normal HSCs were done under self-renewing F12-PVA-based conditions as described previously^22^. Cell-culture-treated 96-well flat-bottom plates were coated with 100 ng/ml fibronectin (Bovine Fibronectin Protein, CF Catalog: 1030-FN) for 30 minutes at room temperature. Following sorting, HSCs were centrifuged 800 g for 8 minutes, resuspended with concentrated LARRY lentiviral particles (typically 20-50 ul, depending on viral titers), and then transferred into 200 μl of complete HSC medium supplemented with 100 ng/ml recombinant mouse TPO and 10 ng/ml recombinant mouse SCF (PeproTech: TPO AF315-14; SCF AF250-03). HSCs were grown at 37°C with 5% CO2. Multiplicity of infection was typically 0.15-0.2, with an estimated 1.2 integrations per cell. When lentiviral library transduction was performed, the first media change took place 24 hours post-transduction. All other protocol steps followed the guidelines provided in ref. 22. After 5-6 days of expansion, all live LARRY-labelled (EGFP^+^) cells were FACS sorted and transferred to PLSTC medium supplemented with 100 ng/ml recombinant mouse TPO, 50 ng/ml recombinant mouse SCF, 10 ng/ml recombinant mouse IL-3, 10 ng/ml recombinant mouse IL-6 (PeproTech: TPO AF315-14; SCF AF250-03; IL-3 AF213-13; IL-6 AF216-16) and grown at 37°C with 5% CO2. From day 5 to day 10 of culture, 4-hydroxitamoxifen (4OHT, previously dissolved in ethanol) was added at a final concentration of 2 μM to induce the leukemic mutations. LSCs were subsequently maintained under the same polymer-based culture conditions: 96-well fibronectin-coated plates, medium change every other day, and splitting when cultures reached ∼80% confluence. Cells were highly sensitive to medium freshness; therefore, aliquots of 10–20 ml were prepared, sealed with Parafilm, and used within a maximum of 2 weeks. Cells could be replated into the same fibronectin-coated well, and we did not notice a difference in phenotype. It is crucial to avoid letting the wells dry after removing the fibronectin coating buffer, as this results in loss of viability. Cells show a marked preference for growing in 96-well plates (compared to 48-well or 24-well formats). Polymer-based cultures in an exponential growth phase can be frozen as described above using 100-μl freezing media per well (approximately 100,000-200,000 cells).

## Construction of lentiviral pLARRYv3/Multi-index EGFP vectors

The construction of barcoded libraries was executed by a previously established protocol (https://www.protocols.io/view/barcode-plasmid-library-cloning-4hggt3w). First, the EGFP coding sequences and the EF1a promoter sequences were PCR amplified from pLARRY-EGFP with primers homologous to the vector insertion site in a custom synthetic lentiviral plasmid backbone (Vectorbuilder, Inc) using Gibson assembly (Gibson Assembly® Master Mix, NEB, Ref. E2611L). After magnetic-bead purification, ligated vectors were then transformed into NEB10-beta electroporation ultracompetent E.coli cells (NEB® 10-beta Electrocompetent E. coli, NEB, Ref.C3020K) and grown overnight on LB plates supplemented with 50 μg/mL Carbenicillin (Carbenicillin disodium salt, Thermo Scientific Chemicals Ref. 11568616). Colonies were scrapped using LB medium and pelleted by centrifugation. Plasmid maxipreps were performed using the Endotoxin-Free Plasmid Maxi Kit (Macheray Nagel), following the manufacturer’s protocol. pLARRY-EGFP was a gift from Fernando Camargo (Addgene plasmid 140025).

## Barcode lentivirus library generation

To barcode pLARRYv3 (Multi-indexed EGFP) plasmids and generate barcoding libraries, first a spacer sequence flanked by EcoRV restriction sites was cloned into the plasmid after the WPRE element of the vector. Custom PAGE-purified single-strand oligonucleotides with a pattern of 20 random bases and surrounded by 25 nucleotides homologous to the vector insertion site were synthesized by IDT DNA Technologies (**Supplementary Table 13 - primers**). The assembly of these components and subsequent purification steps were carried out through Gibson assembly (Gibson Assembly® Master Mix, NEB, Ref. E2611L). Six electroporations of the bead-purified ligations were performed into NEB10-beta E.coli cells (NEB® 10-beta Electrocompetent E. coli, New England Biolabs, Ref.C3020K) utilizing a Gene Pulser electroporator (Biorad). Subsequently, after transformation, the cells were incubated at 37 °C for 1 hour at 220 rpm. Post-incubation, the transformed cells were plated in six large LB-ampicillin agar plates overnight at 30°C. Colonies from all six plates were collected by scraping with LB-ampicillin and then grown for an additional 2h at 225 rpm and 30 °C. Cultures were pelleted by centrifugation, and plasmids were isolated using the Endotoxin-Free Plasmid Maxi Kit (Macheray-Nagel), following the manufacturer’s protocol. Lentivirus production and HSPC transduction were performed as described in^36^. Library diversity was ∼220,000 barcodes per LARRY-index, as estimated by capture-recapture experiments (i.e. number of unique vs number of common sequences from multiple independent transduction experiments).

## In vivo clonal response experiment and therapy modeling

For in vivo experiments, we transplanted EGFP+ leukemic cells (previously expanded in StemSpan or PLSTC conditions) into sub-lethally irradiated mice (9 Gy, split dose of 4.5 Gy). We used CD45.1 mice (Charles River France, B6.SJL-PtprcaPepcb/BoyCrCrl) as recipients. For PLSTC versus Stemspan comparisons, we transplanted 1000, or 100,000 whole-well EGFP+ (leukemic) cells per recipient. In all cases, cells were counted with a LUNA-FL device and resuspended in PBS (100 ul per recipient). For the other transplants (Figure 3 and Figure 4), we used 100,000 whole-well EGFP+ cells per recipient. For *in vivo* treatments (Figure 4), recipient mice were randomly assigned to the two different treatment groups (group 1 and group 2). Treatment typically began on day 14 after transplantation. Group 1 received an induction chemotherapy regimen of 5 days of cytarabine (araC, Sigma-Merck, cat. No. PHR1787, 100 mg kg−1) plus 3 days of doxorubicin (Doxo, Sigma-Merck, cat. No. PHR1789, 3 mg kg−1) delivered by intraperitoneal (i.p.) injection as previously described^17^. Group 2 received i.p. injections of equivalent DMSO (vehicle) volume following the same dosing regimen as for group 1 mice. One week after, therapeutic response was confirmed by peripheral blood sampling. Bone marrow and spleen cells from vehicle mice and treated mice were collected 35 days after transplantation. Mice were euthanised, and bone marrow (femur, tibia, and pelvis) and spleen were harvested through mechanical force. The collected cells were then sieved through a 100-μm strainer and cleansed with a cold ‘Easy Sep’ buffer containing PBS with 2% fetal bovine serum (FBS), followed by lysis of red blood cells using RBC lysis buffer (Biolegend, Catalog no. 420302). These cells were then washed, blocked with FcX, and stained with the following fluorescently labeled antibodies: APC anti-mouse cKit and DAPI. In addition, to multiplex different organs and donors within the same reactions, cells were incubated with cell-hashing Mouse TotalSeq-B antibodies (Biolegend, TotalSeq-B Mouse barcodes 1-8). Barcodes 1 through 4 identified the bone marrow cells for each recipient, and barcodes 5 through 8 identified the spleen cells for each recipient). cKit^+^ and cKit^−^ barcoded (EGFP^+^, T-Sapphire^+^) leukemic cells were sorted pooled in a 1:1 ratio for constructing a single-cell library using Chromium Single Cell 3’ Reagent Kits (v3.1 with Feature Barcoding) following the manufacturer’s guidelines (10X Genomics). Barcode library, feature barcode, and gene expression libraries were prepared as described below.

## Single-cell encapsulation, transcriptome and Feature-barcode library preparation

For scRNA sequencing and subsequent re-plating, cells were pipetted up and down gently a few times to dissociate into single cells and transferred to a 1.5 mL microtube. The well was then washed with 100 ul of EasySep buffer to collect all the possible remaining cells. Cells were then concentrated by centrifugation at 800 g for 8 minutes. Washed cells were then blocked with FcX, and stained with CD34, CD48, CD16/32 and FceRI antibodies. To minimize the impact of batch effects on sequencing, we multiplexed different conditions leveraging the unique barcode pattern of our libraries together with Biolegend TotalSeq™ anti-mouse hashing antibodies, enabling the simultaneous preparation of libraries representing all experimental conditions in a single reaction for each day of sampling. All live cells were sorted based on fluorescent reporter expression depending on each experiment (e.g. LARRY-EGFP). Part of the sample (as specified in the main text) was transferred to a new tube, centrifuged at 800g for 8 minutes, and resuspended in 40 ul of PBS-0.01%BSA for single-cell library preparation. We constructed single-cell whole transcriptome (GEX) and TotalSeq Feature-Barcode (FB) libraries using Chromium Single Cell 3’ Reagent Kits (v3.1) following the manufacturer’s guidelines (10X Genomics). The remaining part was then plated back for further expansion in culture.

## Single-cell library preparation for LARRY sequencing

Following the reverse transcription of mRNA and first-strand cDNA amplification, 100 ng of the cDNA libraries were used as templates to amplify LARRY barcodes by nested PCR similar to the protocol described in Singh et al^16^. The first PCR used forward primer (Pre-Enrichment forward) CTG AGC AAA GAC CCC AAC GAG AA together with the corresponding 10x Genomics dual index TruSeq reverse primer using the following programs 1, 98 C, 3 min; 2, 98 C, 20 s; 3, 58 C, 15 s; 4, 72 C, 20 s; 5, repeat steps 2–4 08 times; 6, 72 C, 3 min; 7, 4 C, hold. The PCR products were then purified with a 0.8:1 ratio of Ampure XP beads. Purified PCR products were then subjected to a second PCR using the forward primer (Trueseq_LARRY) GTG ACT GGA GTT CAG ACG TGT GCT CTT CCG ATC TGC TAG GAG AGA CCA TAT GGG ATC and the corresponding 10x dual index Truseq reserve primer, following program 1, 98 C, 3 min; 2, 98 C, 20 s; 3, 58 C, 15 s; 4, 72 C, 20 s; 5, repeat steps 2–4 08 times; 6, 72 C, 3 min; 7, 4 C, hold. The final PCR products were then purified by a 0.8:1 ratio of Ampure XP bead: PCR products, were indexed using the 10x dual index TruSeq kit, and sequenced using Illumina NovaSeq or NextSeq.

## CROPseq experiments

To select the candidate genes for perturbation-sequencing, we examined positively-enriched differentially-expressed genes in the PLSTCs versus StemSpan single-cell transcriptomics experiment across the LSC-like leiden cluster 2. We then cross-checked this with our reanalysis of primitive versus mature NPM1-mutant AML single-cell data. From the common set of genes, we cherry-picked candidates subjectively based on their primitive AML-specific gene expression and our perceived potential for therapeutic translation using bibliographic search (inhibitors available, receptors that could be targetable by immunotherapy, and reduced expression in critical tissues such as heart, liver and kidney). We then designed sgRNAs using the Broad Institute CRISPick pipeline (library mode) and selected 2 non-targeting controls (NTC1 and NTC2), 2 essential genes (Gapdh, Plk1), and 6 candidates (Adgrg1, Csgalnact1, Mat2a, Spns2, Zeb1 and Mat2a). The final sgRNA library was ordered from Twist Bioscience to be cloned into our custom plasmid (CROPseq-T-Sapphire). Large fragment cloning was performed placing 2 sgRNAs per plasmid, separated by a tRNA-linker sequence (**Supplementary Table 14 - guide RNAs**). After plasmid midi-prep, we used Lenti-X 293 cells to produce CROPseq lentiviral particles. PLSTC-expanded leukemic cells were transduced with an SFFV-SpCas9-mScarlet lentivector and sorted to enrich mScarlet+ primitive LSCs (CD16/32^−^). Multiple batches of primitive LSCs in separate wells were transduced with each CROPseq sgRNA plasmid (4 wells per target). After 8 days in culture, PLSTC cells were stained with TotalSeq Mouse Antibodies (Biolegend, barcodes 1-4), sorted for live mScarlet+ T-Sapphire+ cells, and pooled into a single 10x Chromium X lane using the v3.1 Gene Expression with Feature Barcoding kit for single-cell transcriptome library preparation.

## Single-cell library preparation for CropSeq

For Cropseq sublibrary preparation, 100 ng of the cDNA libraries were used as templates to amplify gRNAs by nested PCR similar to the protocol described above: the first PCR used forward primer (Pre-Enrichment forward) CTG AGC AAA GAC CCC AAC GAG AA together with the corresponding 10x Genomics dual index TruSeq reverse primer using the following programs 1, 98 C, 3 min; 2, 98 C, 20 s; 3, 58 C, 15 s; 4, 72 C, 20 s; 5, repeat steps 2–4 08 times; 6, 72 C, 3 min; 7, 4 C, hold. The PCR products were then purified with a 1,5:1 ratio of Ampure XP beads. Purified PCR products were then subjected to a second PCR using the forward primer (TruseqR2_Cropseq) GTG ACT GGA GTT CAG ACG TGT GCT CTT CCG ATC TCT TGT GGA AAG GAC GAA ACA CCG and the corresponding 10x dual index Truseq reverse primer, following program 1, 98 C, 3 min; 2, 98 C, 20 s; 3, 58 C, 15 s; 4, 72 C, 20 s; 5, repeat steps 2–4 08 times; 6, 72 C, 3 min; 7, 4 C, hold. The final PCR products were then purified by a 1,5:1 ratio of Ampure XP bead: PCR products, were indexed using the 10x dual index TruSeq kit, and sequenced using Illumina NovaSeq or NextSeq^81^.

## Single-cell RNAseq data processing and calling of lineage barcodes (CloneRanger)

Single-cell RNA-seq raw data was processed with CellRanger (v10.0.0). Lineage barcode and CROP-seq guide reference sequences were defined and processed using **CloneRanger (<u>github.com/dfernandezperez/cloneranger</u>)** to (i) generate Hamming-distance–corrected reference libraries for expressed lineage barcodes (LARRY) and CROP-seq sgRNAs, (ii) integrate these custom features into a unified feature reference file, and (iii) enable simultaneous quantification of gene expression, cell hashing oligonucleotides (TotalSeq), lineage barcodes, and sgRNA transcripts. Custom feature reference files produced by CloneRanger were supplied to CellRanger and sequencing data were processed using Cell Ranger with custom features, allowing concurrent mapping of conventional gene expression features together with TotalSeq antibody hashtags, LARRY lineage barcodes (or CROP-seq guide RNA sequences). Feature counts supported by at least 3 reads were maintained. CellRanger outputs consisted of filtered feature-barcode matrices, feature annotation tables, and barcode tables containing counts for all detected transcripts and features. These processed matrices, provided as processed files in GEO, served as the starting point for all downstream computational analyses. Downstream analyses were performed in Python (v3.11) using Scanpy (v1.11.5). Cells were filtered using dataset-specific thresholds on the number of detected genes, total UMI counts, and the fraction of mitochondrial transcripts. Thresholds were selected manually after inspecting data distributions for each library. For multiplexed experiments, TotalSeq hashtag counts were extracted from the feature matrices and sample demultiplexing was performed using `hashsolo` (Scanpy external). Only confidently classified singlet cells were retained, and cells classified as doublets or negatives were excluded from further analysis. We further used a second doublet identification tool, Scrublet, to further identify and exclude potential doublets. LARRY expressed lineage barcodes were identified based on their feature name within the count matrices. LARRY barcode features with fewer than 3 total counts per cell were set to zero to remove background signal. Each cell was assigned a unique lineage barcode by selecting the barcode with the highest remaining count. Cells lacking detectable barcode expression after thresholding were labeled as negative and excluded. LARRY barcode-associated features were removed from the expression matrix after assignment to prevent downstream bias. Similarly, for CROP-seq experiments, sgRNA features were thresholded by requiring at least 3 counts per cell, and each cell was assigned a single sgRNA identity based on maximal sgRNA expression. Cells without detectable sgRNA expression or classified as doublets were excluded. sgRNA features were removed from the expression matrix after guide assignment. Final processed datasets were stored as AnnData h5ad objects and used for all subsequent analyses.

## Single-cell integration, clustering, and annotation (Scanpy/Seurat)

After preprocessing, gene expression matrices were normalized to a total of 10,000 counts per cell and log-transformed. Highly variable genes (HVGs) were selected using the default Scanpy method, retaining the top 2,000 genes based on mean expression and dispersion thresholds. Then, cell-cycle–associated genes were identified using canonical S– and G2/M-phase gene sets and either regressed out or explicitly excluded from the HVG set prior to dimensionality reduction. Principal component analysis (PCA) was performed on the scaled HVG expression matrix, and the number of retained components (typically 30–50 PCs) was chosen based on explained variance. For Figure 1 (PLSTCs versus StemSpan cultures) low-dimensional representations were integrated using Harmony, and Harmony embeddings were used for neighborhood graph construction and visualization. Cell neighborhood graphs were constructed using the k-nearest neighbors (typically k = 15–45, depending on dataset size) in the integrated PCA space. Uniform Manifold Approximation and Projection (UMAP) was used for visualization. Graph-based clustering was performed using the Leiden algorithm with resolution parameters tuned per dataset (typically 0.2–1.5, depending on complexity). Clusters corresponding to low-quality cells or technical artifacts were identified based on quality control metrics (log-size differences in average mitochondrial gene counts or total counts) and removed prior to final clustering. Cell-state annotation was performed by examining the expression of Leiden cluster differentially-expressed genes, curated gene signatures, and bibliography. Marker genes for each cluster were identified using Wilcoxon rank-sum tests as implemented in Scanpy. For day 40 *in vivo* (control versus treated) samples, clustering and embedding were repeated after subsetting all the cells belonging to clusters with positive enrichment for all the LSC signatures.

## Gene signature definition and scoring

Curated gene signatures were defined a priori and used to quantify cell-cycle status, stemness, lineage-associated programs, and transcriptional memory states. All gene signatures were defined explicitly as gene lists in the analysis code and are reported in **Supplementary Tables 4 and 8**. Signature scores were computed using Scanpy (`sc.tl.score_genes`) on normalized and log-transformed expression data. Leukemia stem cell associated transcriptional programs were extracted by taking only the positively-enriched LSC genes from scores developed in Ng et al. 2016^33^. Lineage-associated transcriptional signatures were quantified curating gene lists from our previous analyses^36,38^. Additional custom gene signatures were defined based on the results from this manuscript (e.g. resistance-memory signature).

## Sister cell similarity analysis

To quantify transcriptional similarity between sister cells derived from the same lineage, cells were grouped according to their assigned lineage barcode. Only barcodes represented by at least two cells within a given experimental condition were considered for analysis. Similarity between pairs of cells was computed using cosine similarity (Scipy v1.15.2) between their PC embeddings. Pearson correlation, implemented using SciPy, was used as a secondary metric for validation in selected analyses. For each lineage barcode, all pairwise similarities between sister cells were calculated, and the resulting values were aggregated across all clones. To assess whether observed sister-cell similarity exceeded expectations by chance, a label-permuted control was generated by randomly shuffling lineage barcode assignments while preserving the empirical clone size distribution. Statistical significance was assessed by comparing observed sister-cell similarity distributions to control distributions using Wilcoxon rank-sum tests.

## Fate clustering with scLiTr

Clone-level fate representations were constructed using scLiTr v1.0.1 (github.com/kharchenkolab/scLiTr). scLiTr was used for clone-aware graph construction, clone-level fate vectorization, label transfer between clone– and cell-level objects, clone-level expression summarization, and clone-aware visualization. Clone-aware nearest-neighbor structure was computed using `sl.tl.clonal_nn`, providing the AnnData object with lineage barcode annotation as the clone identifier and requiring a minimum clone size of 3–5 cells, depending on the dataset. Clone-level fate vectors were computed using `sl.tl.clone2vec` by aggregating cell-state annotations within each clone. Specifically, each clone was represented by the relative frequencies of cells across Leiden clusters, yielding a clone-by-fate representation. Clone embeddings were constructed from the clone-level fate vectors and used to build clone–clone neighborhood graphs (5 nearest neighbors), followed by UMAP visualization and Leiden clustering (resolutions 0.4-0.8, depending on complexity). Clone cluster assignments were transferred back to individual cells using `sl.tl.transfer_clonal_annotation`, enabling visualization and stratification of cells by clone-level fate class. Clone-aware visualization of cell embeddings was performed using scLiTr plotting utilities, such as density-based visualizations using `sl.pl.kde` (Kernel density bandwidths were set explicitly, 0.09–0.2, depending on the number of cells per clonal group). To enable clone-level transcriptional analyses, clone-aggregated expression profiles were computed using `sl.tl.summarize_expression`. The resulting clone-level AnnData objects were used for gene expression visualization, gene signature scoring, and differential expression analyses using Scanpy.

## Stochastic sampling model of clonal selection and fate bias

Stochastic sampling models and permutation-based tests were used to quantify clonal selection, fate bias, and engraftment probabilities across experimental contexts. Clone-level fate class labels (e.g., scLiTr-derived clone clusters) and clone-level summary metrics (such as the fraction of cells from a given clone detected in a target context) were extracted and treated as categorical or continuous variables, respectively. To assess whether specific clone classes exhibited non-random enrichment or depletion in a given context (for example, preferential engraftment of clones belonging to a particular fate class), observed statistics were computed from the data, such as the fraction of clones or cells of a given class present in the target condition, or the distribution of clone contributions across conditions. Null distributions were generated by permuting clone labels. Permutations preserved clone size distributions and, where appropriate, stratification by replicate (leveraging the TotalSeq hashing). For comparisons between matched contexts (e.g., D20 ex vivo versus D20 in vivo), permutations were restricted within matched strata, typically performing label shuffling in one of the two groups. In most cases 5,000 permutations were performed per test. Effect sizes were quantified as differences or ratios between observed values and the mean of the null distribution. Empirical p-values were computed. When multiple classes were tested in parallel, p-values were adjusted for multiple testing using the Benjamini-Hochberg FDR procedure.

## Single-cell differential gene expression

Differential gene expression analyses were primarily performed using Scanpy on normalized and log-transformed expression data. Unless otherwise stated, differential expression was assessed using the Wilcoxon rank-sum test as implemented in <u>`sc.tl`</u>`.rank_genes_groups`. For clone-level differential expression, clone-aggregated expression profiles generated using `sl.tl.summarize_expression` were analyzed using scLiTr package. Differential expression between scLiTr-defined clone classes was performed using `sc.tl.rank_genes_groups` on clone-level AnnData objects. For comparison of PLSTIC and Stemspan conditions (Fig. 1), pseudobulk differential expression was performed using DESeq2.

## Pathway Analysis and Transcription Factor Activity Predictions

Pathway and transcription factor (TF) activity inference was performed using Decoupler^54^ by applying a univariate linear model (ULM) to gene expression data. Decoupler analyses were performed on both single-cell and clone-aggregated expression profiles, depending on the specific analysis. For pathway activity inference, PROGENy gene sets were used^55^. For transcription factor activity inference, CollecTRI/DoRothEA-derived TF–target regulatory networks were used, as provided by Decoupler^56^. In additional analyses, MSigDB Hallmark gene sets were used to score pathway activity at the single-cell level. All Decoupler analyses were performed using ULM via `dc.mt.ulm`, with a minimum target gene threshold of tmin = 3-4, depending on the resource. Differential activity across experimental conditions, cell states, clone classes, or fate annotations was assessed using Wilcoxon rank-sum tests, using `dc.tl.rankby_group`.

## Single cell Perturbation and Transcriptome Sequencing analysis

For CROP-seq perturbation enrichment analyses, sgRNA representation across cellular states was evaluated using contingency-table methods. Enrichment of individual sgRNAs within Leiden clusters was tested using Fisher’s exact tests (2×2 tables: sgRNA vs not-sgRNA, in-cluster vs out-of-cluster), with p-values adjusted using Benjamini-Hochberg FDR. To assess sgRNA enrichment while accounting for replicate structure, we used Cochran-Mantel-Haenszel (CMH) tests using `statsmodels.stats.contingency_tables.StratifiedTable`, stratifying by replicate. CMH pooled odds ratios and p-values were computed per sgRNA relative to background, with FDR correction across sgRNAs. Neighborhood-level differential abundance in the CROP-seq dataset was assessed using Milo, implemented in pertpy (v1.0.2)^111^. kNN graphs were constructed with 100 neighbors on PCA space, neighborhoods were defined with `milo.make_nhoods`, and neighborhood counts were summarized by sample. Differential abundance testing was performed with `milo.da_nhoods` using a DESeq2-style negative binomial model via `pydeseq2`, and significance was evaluated using SpatialFDR. Neighborhood graphs and DA summaries were visualized using `milo.plot_nhood_graph` and `milo.plot_da_beeswarm` with explicit SpatialFDR thresholds (alpha = 0.1) and minimum neighborhood sizes (min_size = 5-10).

## Statistical methods

For single-cell data, statistical analyses were performed using Python (v3.11) with NumPy (v2.3.3), SciPy (v1.15.3), pandas (v2.3.3), and Scanpy (v1.11.4), unless otherwise stated. For flow cytometry, survival curves, and other cell phenotyping data, statistical analyses were performed using Prism version 10, typically using two-way ANOVA (time series data analysis) or Mann-Whitney rank tests. All statistical tests were two-sided and non-parametric, and no assumptions of normality were made unless explicitly noted. For comparisons of gene expression, gene-signature scores, pathway activity scores, and transcription factor activity scores between groups, Wilcoxon rank-sum tests were used. For categorical enrichment analyses (e.g., overrepresentation of clone classes across conditions or clusters), Fisher’s exact tests were used. For permutation-based statistical tests, empirical p-values were computed. Multiple-testing adjustment of p-values used the Benjamini-Hochberg FDR procedure. Statistical significance thresholds are indicated in the corresponding figure legends, with asterisks marking the threshold reached.

